# Pharmacokinetic considerations for optimizing inhaled spray-dried pyrazinoic acid formulations

**DOI:** 10.1101/2023.04.01.534965

**Authors:** Shekhar B. Yeshwante, Patrick Hanafin, Brittany K. Miller, Laura Rank, Sebastian Murcia, Christian Xander, Ayano Annis, Victoria K. Baxter, Elizabeth J. Anderson, Brian Jermain, Robyn Konicki, Alan A. Schmalstig, Ian Stewart, Miriam Braunstein, Anthony J. Hickey, Gauri G. Rao

## Abstract

Tuberculosis (TB), caused by *Mycobacterium tuberculosis* (*Mtb*), remains a leading cause of death with 1.6 million deaths worldwide reported in 2021. Oral pyrazinamide (PZA) is an integral part of anti-TB regimens, but its prolonged use has the potential to drive development of PZA resistant *Mtb*. PZA is converted to the active moiety pyrazinoic acid (POA) by the *Mtb* pyrazinamidase encoded by *pncA*, and mutations in *pncA* are associated with the majority of PZA resistance. Conventional oral and parenteral therapies may result in subtherapeutic exposure in the lung, hence direct pulmonary administration of POA may provide an approach to rescue PZA efficacy for treating *pncA-*mutant PZA-resistant *Mtb*.

The objectives of the current study were to i) develop novel dry powder POA formulations ii) assess their feasibility for pulmonary delivery using physicochemical characterization, iii) evaluate their pharmacokinetics (PK) in the guinea pig model and iv) develop a mechanism based pharmacokinetic model (MBM) using *in vivo* PK data to select a formulation providing adequate exposure in epithelial lining fluid (ELF) and lung tissue.

We developed three POA formulations for pulmonary delivery and characterized their PK in plasma, ELF, and lung tissue following passive inhalation in guinea pigs. Additionally, the PK of POA following oral, intravenous and intratracheal administration was characterized in guinea pigs. The MBM was used to simultaneously model PK data following administration of POA and its formulations via the different routes. The MBM described POA PK well in plasma, ELF and lung tissue.

Physicochemical analyses and MBM predictions suggested that POA maltodextrin was the best among the three formulations and an excellent candidate for further development as it has: (i) the highest ELF-to-plasma exposure ratio (203) and lung tissue-to-plasma exposure ratio (30.4) compared with POA maltodextrin and leucine (75.7/16.2) and POA leucine salt (64.2/19.3); (ii) the highest concentration in ELF (*Cmac_ELF_*: 171 nM) within 15.5 minutes, correlating with a fast transfer into ELF after pulmonary administration (*k_PM_*: 22.6 1/h).

The data from the guinea pig allowed scaling, using the MBM to a human dose of POA maltodextrin powder demonstrating the potential feasibility of an inhaled product.

**Table of Contents (TOC)/Abstract Graphic:** 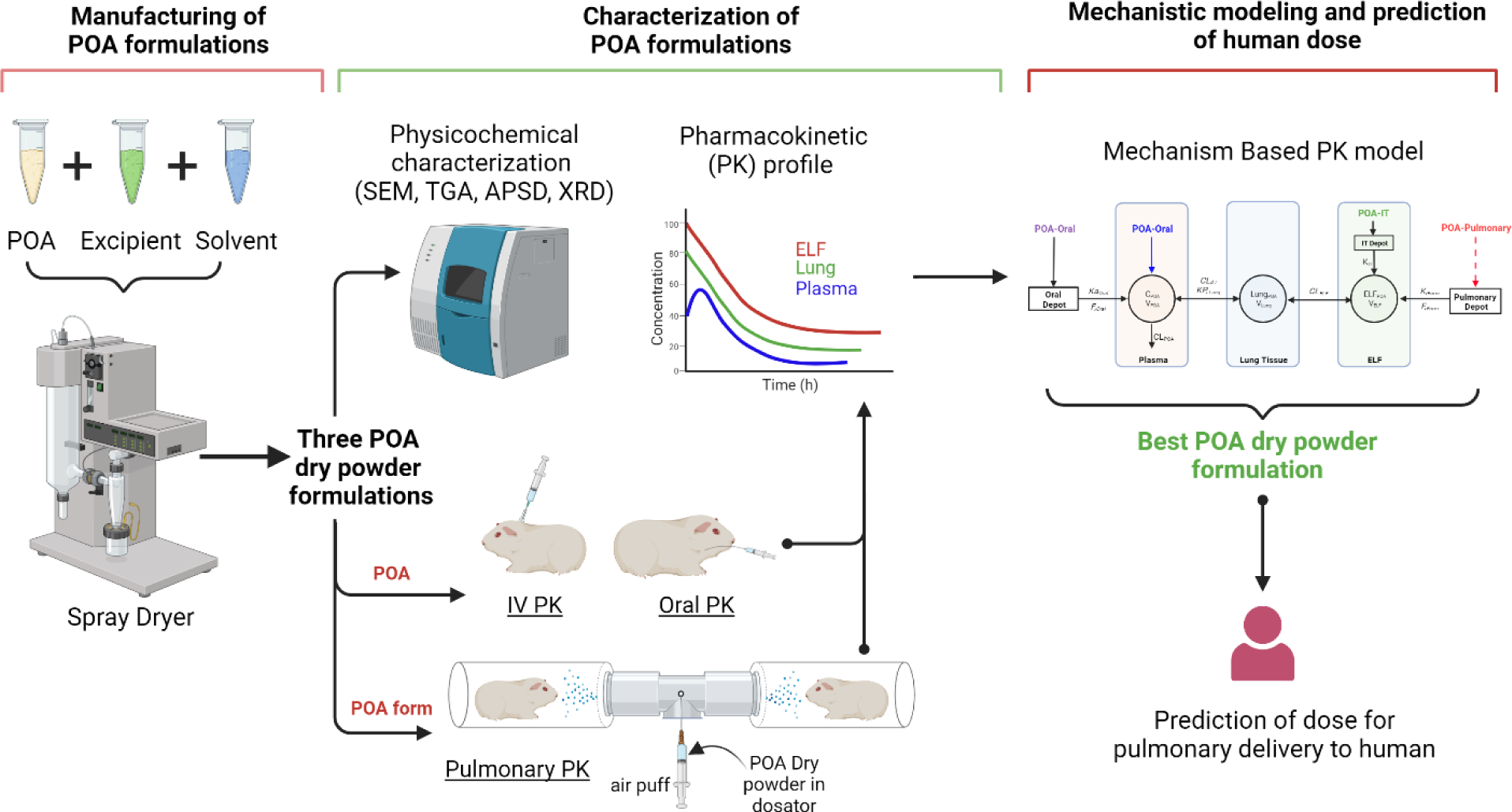

## INTRODUCTION

Tuberculosis (TB), caused by the bacterial pathogen *Mycobacterium tuberculosis (Mtb)*, currently ranks second only to SARS-CoV-2 as the deadliest infectious disease. In 2021, there were 10.6 million new *Mtb* infections and 1.6 million *Mtb*-related deaths worldwide, marking the first time in over a decade that deaths from TB increased.^1^ Further, one-fourth of the global population are thought to be infected with *Mtb*.^2^ Compounding the impact of *Mtb*, multi-drug resistant TB (MDR-TB) cases are increasing, significantly challenging ongoing efforts to end the TB crisis.^1^

TB is usually treated with complicated multi-drug therapy regimens over an extended duration. Patients infected with drug-susceptible TB (DS-TB) are treated with a four drug combination and, depending on the regimen, treatment duration can range from 4-6 months.^3^ MDR-TB treatment is based on the susceptibility profile of the clinical isolate and can last from six months to over two years.^4, 5^ Treating MDR-TB is further challenged by lack of adequate drug exposure at the site of infection, the prevalence of adverse effects associated with many second-line TB drugs, and a lack of novel treatment strategies effective against MDR-TB. Along with new drugs, strategies to shorten duration of therapy, improve adherence, and limit periods of transmission are needed. Given this need, optimizing approved TB drugs to ensure efficacious drug exposures in lung tissue and epithelial lining fluid (ELF) is essential to limit the development of resistance and deserves consideration.^6, 7^

Pyrazinamide (PZA), a prodrug, is a common and critical component of multi-drug regimens for both DS-TB and MDR-TB.^8, 9^ It is metabolized to the active moiety pyrazinoic acid (POA) by *Mtb*. PZA is distinguished from many other TB drugs by its sterilizing activity and ability to kill *Mtb* while it is in a slow-growing and non-replicating persister state.^10, 11^ PZA is also active in the acidic, nutrient-starved, and hypoxic environments associated with the multi-cellular granulomas that typify the host immune response to *Mtb*.^10^ Importantly, PZA can penetrate cellular and necrotic granulomas, which does not occur for all TB drugs.^12^ Finally, PZA has synergistic activity in combination with other anti-*Mtb* drugs.^11^ For this reason, PZA is often included when new TB drug regimens are studied (*e.g*., BPaMZ, endTB, MDR-END, NeXT, TRUNCATE-TB).^8, 13–15^

However, PZA’s role as a cornerstone TB treatment is challenged by the rise in PZA-resistant *Mtb*. Indeed, 50% of MDR *Mtb* strains are PZA resistant.^16, 17^ PZA is converted to POA by the *Mtb* pyrazinamidase (PZAse) enzyme encoded by the *pncA* gene.^18, 19^ Additionally, host-mediated enzymatic conversion of PZA to POA has been reported in mice, guinea pigs, and humans.^20^ However, the largest category of PZA-resistant *Mtb* mutations map to *pncA*, indicating that the majority of POA is not produced by the host.^9, 20^ As the active agent, direct administration of POA has been proposed as a strategy to rescue the PZA efficacy lost in *pncA*-mutant PZA-resistant *Mtb*.^20–22^ Several studies have now shown that POA is active against PZAase-negative *Mtb* strains, including some reporting POA equal to or more, effective than PZA.^10, 22^ However, POA delivered orally was ineffective compared with PZA at equivalent doses in mice^21^ and guinea pigs (unpublished data, Braunstein and Hickey). Pharmacokinetic (PK) studies in mice demonstrate only limited penetration of oral POA into lung ELF compared with PZA, which may explain the poor efficacy observed in animal studies.^21^ We hypothesized that direct pulmonary delivery of POA would overcome this limitation by achieving higher, ideally therapeutic, local POA exposure in the ELF and in lung tissue.

Conventional therapies for respiratory infections delivered orally or parenterally have poor pulmonary distribution. This can result in subtherapeutic drug exposure in the lungs and high systemic concentrations associated with off-target toxicities. Direct pulmonary delivery circumvents the issue of suboptimal exposure within the lung while avoiding first pass metabolism and minimizing systemic exposure.^23, 24^ Pulmonary delivery would be included in a combination regimen where other drugs would treat the systemic manifestations of disease. More recently, studies on the pulmonary delivery of TB drugs to *Mtb*-infected animals indicated the feasibility of inhaled therapy.^25–27^ Understanding drug disposition following pulmonary delivery is important, as exposure at the site of infection determines efficacy. Hence, we evaluated the PK of POA following oropharyngeal inhalation as a strategy to achieve high local POA exposure in the ELF and lung tissue.^23^

Spray dried powder preparations of antimicrobial agents in combination with inhalers that are actuated by the breath of the patient have the advantage of allowing high dose delivery, ease of use, storage and transport stability suitable for use in a wide variety of global environments.^28^ We have demonstrated the feasibility of this approach for several antitubercular drugs, most notably capreomycin sulfate.^29–32^

The objectives of the current study were to 1) develop novel dry power POA formulations suitable for pulmonary delivery; 2) evaluate the PK of these POA formulations in guinea pigs to assess POA exposure in plasma, ELF, and lung tissue; and 3) develop a mechanism-based PK model to characterize POA exposure in plasma, ELF, and lung tissue for optimizing POA formulations with a view to maximizing efficacy.

## MATERIALS AND METHODS

### Drugs and chemicals

Pyrazinoic acid (POA), maltodextrin and reagent-grade ethanol were obtained from Sigma-Aldrich (St. Louis, MO, US). L-leucine was obtained from Alfa Aesar (Ward Hill, MA, USA). KetaVed^®^ (ketamine hydrochloride injection), AnaSed^®^ (xylazine hydrochloride injection), and sodium pentobarbital were from Vedco Inc. (Covetrus, Portland, MA, US), Akorn Animal Health Inc. (Lake Forest, IL, US), and Henry Schein Animal Health (Covetrus), respectively.

### Formulation of POA maltodextrin (PM), POA maltodextrin and leucine (PML), and POA leucine Salt (PLS)

#### Manufacturing of PM, PML, and PLS

A spray dryer (Model B 290, Buchi Corporation, DE, US) equipped with a high efficiency cyclone and two-fluid nozzle (inner orifice and outer orifice 0.7 mm and 1.5 mm, respectively)^33^ was used. POA alone, with excipient-PM, was dissolved in 100% ultrapure water (for each of the POA formulations, 5 mg/mL w/v solids). PML was prepared in 4:1 water: EtOH (v/v) (10 mg/mL w/v solids). The resulting solutions were fed via peristaltic pump through a heated nozzle, where the solutions were atomized using N_2_, resulting in rapid evaporation of the solvent. The remaining dry powder particles were collected downstream via cyclone separator and stored in amber vials at room temperature with a desiccant. In addition to the use of leucine as an additive in the spray-dried product (PML), stoichiometric leucine salt (PLS) was synthesized through the reaction of pyrazinecarboxylic acid (POA) with leucine heated to 60°C, as previously reported.^34^ Yields were calculated as a ratio of powder collected with respect to known starting solid material dispersed in spray-dried solution. Additional details can be found in the **Supplementary Methods**.

#### Physicochemical characterization of particles

Particle morphology was assessed by scanning electron microscopy (SEM; Quanta 200 SEM, FEI, Hillsborough, OR, US). Thermal analysis and moisture content were determined by thermogravimetric analysis (TGA; Q50 TGA, TA Instruments, New Castle, DE, US). Aerodynamic characterization was performed using a Next Generation Impactor (NGI; MSP Corp., Shoreview, MN, US). Particles were characterized for crystallinity by X-ray powder diffraction (XRPD Model D8 instrument, Bruker AXS Inc., Karlsruhe, Germany). Additional details can be found in the **Supplementary Methods**.

#### Aerodynamic characterization

Inertial impaction was performed using an NGI to generate aerodynamic particle size distribution (APSD) following United States Pharmacopeia (USP) <601> procedure outlined for dry powder manufacturing.^35^ A nominal powder mass of 10 mg (including excipients) was loaded into a #3 hydroxypropyl methylcellulose (HPMC) capsule and placed in a RS00 Mod.8 inhaler (Plastiape, Osnago, Italy). The inhaler was actuated by piercing the capsule and then placed in a mouthpiece adapter at the NGI inlet, following which a vacuum was applied. The deposited mass within the NGI aerosol sampling stages and inlet, the inhaler, and capsule were collected by washing with a known volume of deionized water. POA content in the collected samples was quantified by UV spectroscopy (UV-Vis Spectrometer, SynergyMX, Biotech, Winooski, VT, US) at 269 nm. The APSD was reconstructed for the mass deposited at every calibrated NGI stage. The mass median aerodynamic diameter (MMAD), representing the median of the distribution, and the geometric standard deviation (GSD), the measure of the range of particle sizes in the distribution, were derived from plots of NGI data. For a detailed description refer to **Supplementary Methods**.

#### Fine particle dose and fraction

The fine particle dose and fraction (*FPD and FPF* <4.46 μm) were estimated the latter is presented as a function of nominal dose. Specifically, the *FPF* is the ratio of mass collected at impactor stage 3 through the micro-orifice collector relative to the nominal dose (dose in the capsule) and is expressed as a percentage of the nominal dose.

### *In vivo* PK studies

The Institutional Animal Care and Use Committee (IACUC) of the University of North Carolina approved the experimental protocol. For PK characterization, male Dunkin-Hartley guinea pigs (417 ± 62.9 g) with indwelling jugular vein catheters (JVC) were purchased from Charles River Laboratories (Raleigh, NC, US). Guinea pigs were housed individually under standard environmental conditions (ambient temperature 21°C, humidity 60%, 12:12 h light/dark cycle) with food and water *ad libitum*. Guinea pigs were euthanized at the end of each study. Plasma, bronchoalveolar lavage fluid (BALF), and lung tissue were collected and stored at −80°C until LC-MS/MS analysis.

#### Intravenous (IV) PK studies

POA formulated in 0.9% sodium chloride (saline) was intravenously administered to guinea pigs (n=8) via JVCs. Following administration of this single 2 mg/kg IV bolus dose, 0.4 mL of blood was collected at 0.25, 0.5, 0.75, 1, and 1.5 h in lithium heparin tubes (BD Microtainer™ Capillary Collector; BD, Franklin Lakes, NJ, US). Plasma was separated by centrifugation and stored at −80°C until LC-MS/MS analysis.

#### Intratracheal (IT) PK

Tracheostomies were performed in anesthetized guinea pigs (n=4). An 18 G intravenous catheter was inserted into the tracheal lumen and PM dry powder (2 mg/kg) was insufflated through the catheter into the trachea followed by flushing with 10 mL of room air three times. The catheter was removed from the trachea and the incision was closed with a wound clip. Guinea pigs were maintained on 100% oxygen and monitored to ensure sufficient peripheral blood oxygenation. Following intratracheal (IT) administration, 0.4 mL of blood was collected at 0.25, 0.5, 0.75, 1, and 1.5 h via the JVCs.

#### Oral (PO) PK

Guinea pigs (n=9) were administered doses of 300 mg/kg as a suspension in 20% pumpkin puree/ 50% sucrose mixture supplemented with commercial *Lactobacillus* (BD Lactinex) and vitamin C orally. The oral POA dose matched the human-equivalent PZA dose as previously determined by PK analysis in guinea pigs.^20, 36^ Blood (0.4 mL) was collected at 0.08, 0.25, 0.5, 0.75, 1, 1.5, 2, 4, and 6 h via JVCs after oral administration of POA.

#### Pulmonary PK

A custom-built plexiglass, nose-only guinea pig aerosol dosing chamber (**Supplementary Figure 1A**) was used for pulmonary POA PK studies. Twenty-nine guinea pigs (PM (n=11), PML (n=10), and PLS (n=8)) were used. Two guinea pigs were simultaneously dosed with 10 mg of spray-dried PM or PML or 8 mg PLS administered 8 times every 3 mins (**Figure 1**). The formulation was pre-loaded in the dosator and injected into the center of the chamber through an inlet (**Supplementary Figure 1B**). Every spray of the formulation was followed by two puffs of room air (10 mL) every minute to aerosolize the powder. The animals were allowed to inhale the dose for 3 minutes before the subsequent dose was administered. The 3 minute dosing interval was calculated based on the chamber saturation time (2.6 minutes) estimated using the MicroPEM device (RTI MicroPEM, NC, US).^33^ The actual dose inhaled by the guinea pigs was computed using a previously published equation ^37^ representing approximately 3.33 ± 0.098 mg/kg (PM), 1.67 ± 0.028 mg/kg (PML), and 1.65 ± 0.038 mg/kg (PLS) (**see Supplementary Methods**). Blood samples (0.4 mL) were collected at 0.08, 0.17, 0.25, 0.5, 0.75, 1, and 1.5 h via JVCs after administration of all 8 doses every 3 min over 24 mins.

**Figure 1.**
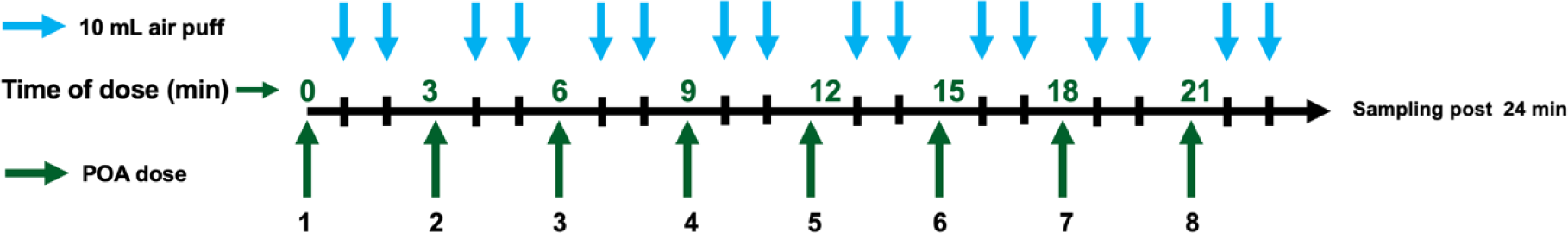
Dosing scheme for pulmonary administration of POA formulations. POA dry powder for direct pulmonary delivery was administered every 3 minutes as indicated by the dark green arrows. Guinea pigs received a total of 8 doses administered every 3 minutes over 24 minutes. An air puff (10 mL) was administered every minute as indicated by the blue arrows, twice after each dose administration, to assist with aerosolization of the powder deposited in the chamber. Blood samples for quantification of POA in plasma were obtained after the last dose was administered (i.e., after 24 minutes).

### Bioanalytical assay for POA quantification

POA was quantified using an LC-MS/MS system consisting of AB-Sciex API 3200 triple-quadrupole mass spectrometer (AB SCIEX, Foster City, CA, US) with an Agilent 1100 HPLC system (Agilent Technologies, Santa Clara, CA, US). Agilent Eclipse XDB (C_8_, 150 x 4.6 mm, 5 µM) was used as the analytical column. A mobile phase of 0.1% formic acid in water (A) and acetonitrile (B) was used at a flow rate of 1 mL/min with a gradient elution starting from 20% B. The mass transitions (Q1/Q3) of *m/z* 125.216/106.786 and 609.263/194.9 were used for POA and reserpine (internal standard), respectively. The lower limit of quantification was 202 nM.

#### Quantification of POA in epithelial lining fluid (ELF)

Urea in the plasma and BALF was detected using LC-MS/MS and the QuantiChrom™ Urea Assay Kit (detection limit 13 µM) (BioAssay Systems, Hayward, CA, US). Equation (1) was used to calculate the volume of the ELF (*V_ELF_*), based on urea concentrations measured in the BALF and plasma. The POA concentration in the ELF (*C_POA,ELF_*) was calculated based on POA quantified in BALF (*C_POA,BALF_*) and *V_ELF_* using the relationship in equation (2):^38^

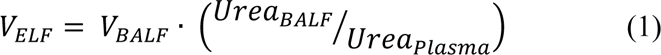

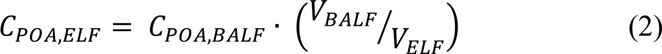

Five mL PBS was used to collect samples from excised lungs resulting in *V_BALF_*, the volume of BALF collected. *Urea_BALF_* and *Urea_Plasma_* are urea concentrations measured in BALF and plasma, respectively.

### PK analysis

#### Non-compartmental analysis

Non-compartmental analysis (NCA) was performed using Phoenix^®^ WinNonlin^®^ version 8.2.0 (Certara USA, Inc., Princeton, NJ, US). The area under the POA plasma concentration vs. time curve up to 1.5 h after dosing (AUC_0–1.5h_) and total AUC were calculated using the linear up - log down trapezoidal rule, and the elimination half-life (t_1/2_), time to reach peak concentration (T_max_), and peak concentration (C_max_) were also determined.

#### Mechanism-based PK model development

Time course PK data obtained following POA administration via the IV, PO, IT, or direct pulmonary routes were simultaneously modeled using a naïve pooled approach with the maximum likelihood estimation (ML) method in ADAPT5 (Biomedical Simulations Resource, Los Angeles, CA, US). Data were visualized using *ggplot2* package (v3.3.6) in R (v4.2.1). The mechanism-based model was developed in a stepwise manner, modeling the time course of POA PK in plasma following IV administration first. This model was then expanded to include oral and intratracheal POA plasma PK data to estimate the first-order oral absorption rate (*Ka_Oral_*) and oral bioavailability (*F_Oral_*) as well as first-order transfer rate (*k_IT_*) between intratracheal depot and ELF and intratracheal bioavailability (*F_IT_*). The model was then expanded by including time course *in vivo* PK for POA formulations following direct pulmonary administration. Formulation-specific parameters, first-order transfer rate (*k_Form_*) between inhalation depot and ELF, and bioavailability after pulmonary administration (*F_Form_*) were estimated.

Linear and nonlinear transfer between the lung tissue and plasma compartments were evaluated. The final model was selected based on model discrimination, evaluating the goodness of fit plots, the precision of the estimated parameters, and AIC values. Model-predicted plasma, ELF, and lung tissue concentration over time profiles were analyzed to estimate *AUC*_0−1.5_*_h_*, *C_max_*, and *T_max_* and these parameters were used to compare different POA formulations.

#### Drug penetration in ELF and lung tissue

Drug penetration into the ELF and lung tissue was calculated as a ratio of model-predicted exposure achieved in the ELF or lung tissue (*AUC*_0−1.5*h,ELF* *or* *lung*_) to the systemic exposure (*AUC*_0−1.5*h,Plasma*_) to assess and compare the penetration of the various formulations into the ELF as well as in the lung tissue.

#### Simulation of human dose

The mechanism-based model was used to estimate the potential first-in-human dose for PM for pulmonary delivery. Simulations were performed using ADAPT5 (Biomedical Simulations Resource, Los Angeles, CA, US) using the “individual simulation with output error” method. Given the lack of information on the PK of POA in humans, we used clinical studies where PZA was administered and the PK of both PZA and POA were characterized to estimate human clearance and volume of distribution. From these clinical studies we considered average clearance and volume of distribution values for POA.^39–41^ Allometric scaling was performed using fixed body weights for humans (BW_H_) and guinea pig (BW_GP_) of 70 kg and 0.42 kg, respectively, to scale other PK parameters from guinea pig (P_GP_) to humans (P_H_) using the equation below:^42^

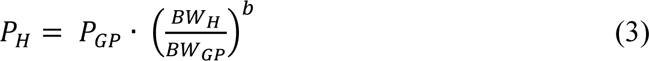

where the values for exponent *b* for volume of distribution, clearance, and first-order transfer rate constants were fixed to 1, 0.75, and −0.25, respectively.^43^ The partition of POA from lung to plasma in humans was assumed to be equal to the guinea pig. The PK/PD index for PZA is *AUC_Plasma_*/MIC of 9.68-14.5 ^44–47^ based on the PZA exposure achieved in plasma. This is considered to be a good predictor of PZA efficacy in the lung considering a 17.8 – 22-fold penetration of PZA to ELF^47, 48^ and C_max_ (>35 mg/L)^45^ is another predictor of efficacy.

Using the scaled PK parameters POA exposure for 1000 subjects were simulated using different doses and dosing frequencies to select the POA dose and dosing frequency. The dose that achieved PK/PD index between 9.68-14.5 was selected as potential human dose for pulmonary administration.

## RESULTS

### Manufactured POA dry powder formulations

Total yield for POA, PM, PML, and PLS dry powders were 60%, 56.7 ± 3.9%, 70.5 ± 2.7%, and 52.2 ± 10.9%, respectively, relative to the initial pre-sprayed mass. X-ray powder diffraction (XRPD) demonstrated that POA, PLS, and PML exhibited crystallinity, the latter showing slight amorphous qualities likely due to the maltodextrin content, which was amorphous as-received (see Supplementary Results).

### Particle morphology and physicochemical characterization

#### Morphology and thermogravimetric analysis

**Figure 2** shows the morphology and physicochemical characterization of spray-dried POA formulations. Without the addition of excipients, POA microparticles formed irregular spheres (**Figure 2A**). However, PM, PML, and PLS formed regular spheres (**Figure 2B-D**). Thermogravimetric analysis in **Figure 2E** indicated that both the precursor POA (black), spray-dried POA (purple), and PLS (green) contained negligible moisture based on negligible mass lost up to 150°C and degradation above 150°C. In contrast, PM (blue) and PML (red) spray-dried samples contained a small amount of moisture (2-3%), likely from the maltodextrin content (dark green).

**Figure 2.**
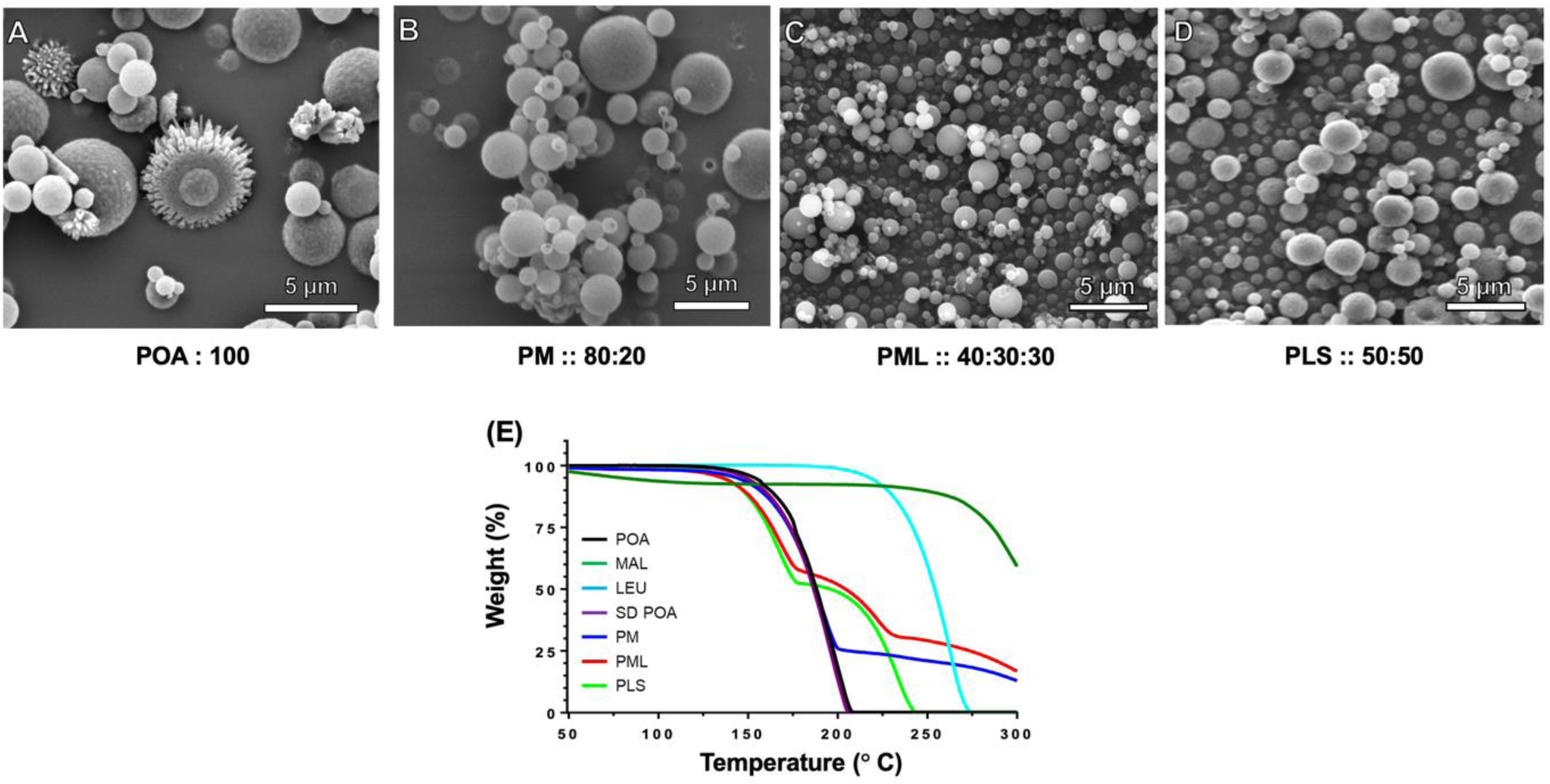
**(A-E)** Results of scanning electron microscopy (SEM) and thermogravimetric (TGA) analysis. SEM images showing the morphology of the spray-dried powder of **(A)** POA, **(B)** PM, **(C)** PML, and **(D)** PLS. **(E)** TGA thermograms for the different POA forms and excipients, pre-sprayed POA (black), pre-sprayed MAL (dark green), pre-sprayed LEU (aqua blue), SD POA (purple), PM (blue), PML (red), and PLS (green). MAL: maltodextrin, LEU: leucine, SD: spray-dried

A semi-quantitative indication of precursor weight content in the spray-dried product of these formulations was assessed by comparing degradation pattern/onset temperatures with their respective starting materials. For example, PML was expected to have 4:3:3 mass ratio and indeed, at ∼59% weight, degradation due to leucine began, followed by maltodextrin at ∼31% (dark green). The conclusions were similar for PM, where the mass ratio was expected to be 8:2 and degradation due to maltodextrin began at ∼25%, and for PLS, where the ratio was expected to be 5:5 and degradation due to leucine began ∼50%.

#### Aerodynamic characterization of POA inhalational formulations

The aerodynamic properties of MMAD, GSD, FPD, and FPF for POA, PM, PML, and PLS are reported in **Table 1**. The POA MMAD of 4.30 µm was large, considering that respirable particles range from 1-5 µm and that 50% of the nominal 10 mg POA dose remained in the capsule or was deposited in the inlet of the NGI, implying that most of this POA aerosol would not enter the lungs. However, PM, PML, and PLS had acceptable MMADs between 2.4 and 3.1 µm. The distributions for PML and PLS were narrower (GSD ∼1.7−1.8) compared with POA and PM (GSD ∼2.0−2.2), indicating a higher likelihood of being within the respirable range. There was an improvement in the FPF of the formulation versus POA, indicating that the proportion of aerosol likely to enter the lungs was much higher: the FPF for POA was ∼15%, while that for PM was ∼68%. However, the highest FPFs were observed for PML and PLS (∼84−86%). Graphical depictions of the APSDs are shown in **Supplementary Figure 2**. XRPD demonstrated that POA, leucine, and PLM exhibited crystallinity, and spray-dried PLS was previously shown to be amorphous (**Supplementary Figure 3**).

**Table 1.**
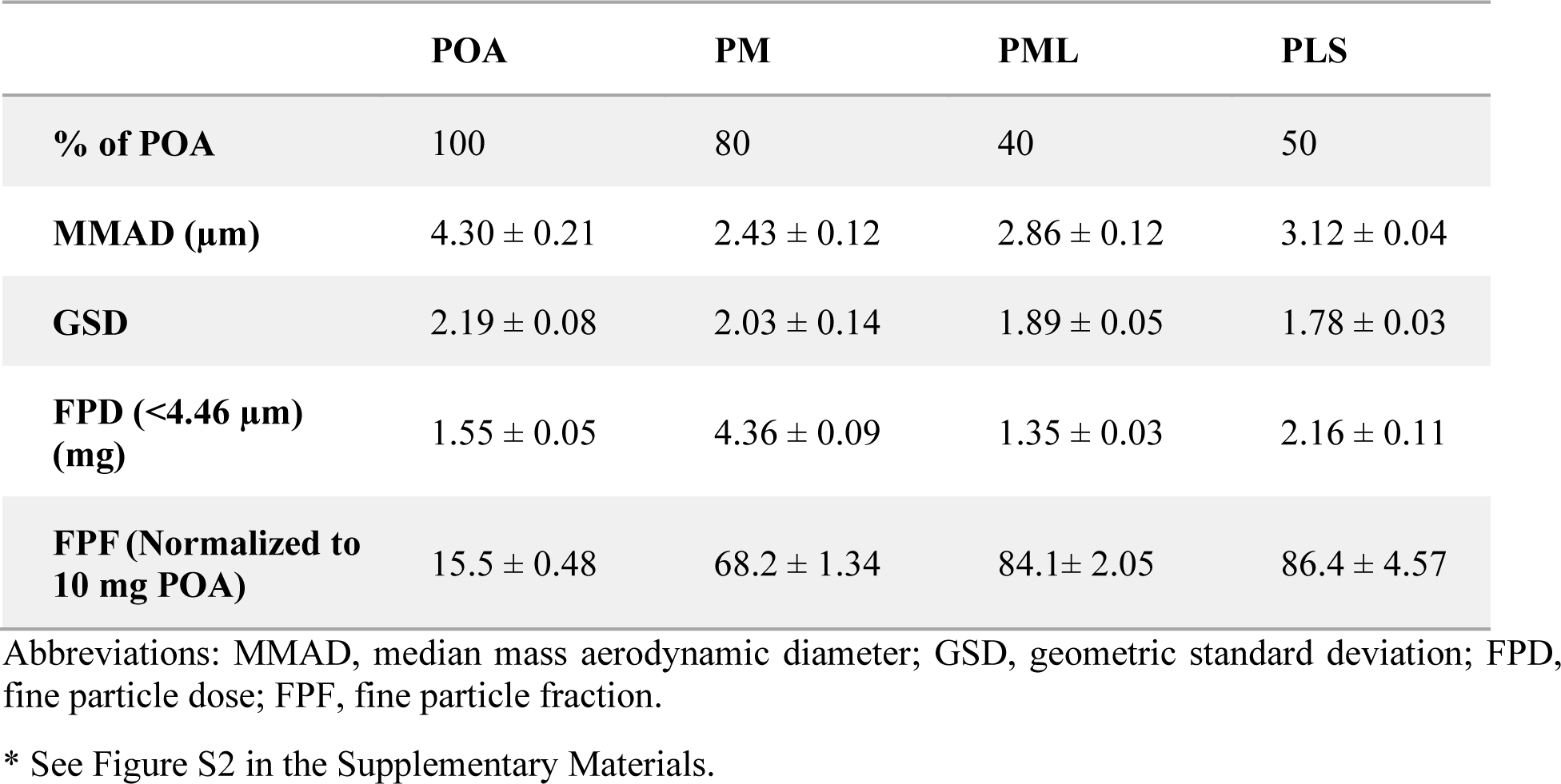
Physiochemical analysis of POA and POA-based formulations based on aerodynamic particle size distribution (APSD)* reported as mean ± SD.

|  | POA | PM | PML | PLS |
| --- | --- | --- | --- | --- |
| **% of POA** | 100 | 80 | 40 | 50 |
| **MMAD ($\mu\text{m}$)** | 4.30 ± 0.21 | 2.43 ± 0.12 | 2.86 ± 0.12 | 3.12 ± 0.04 |
| **GSD** | 2.19 ± 0.08 | 2.03 ± 0.14 | 1.89 ± 0.05 | 1.78 ± 0.03 |
| **FPD (<4.46 $\mu\text{m}$) (mg)** | 1.55 ± 0.05 | 4.36 ± 0.09 | 1.35 ± 0.03 | 2.16 ± 0.11 |
| **FPF (Normalized to 10 mg POA)** | 15.5 ± 0.48 | 68.2 ± 1.34 | 84.1 ± 2.05 | 86.4 ± 4.57 |

Abbreviations: MMAD, median mass aerodynamic diameter; GSD, geometric standard deviation; FPD, fine particle dose; FPF, fine particle fraction.
\* See Figure S2 in the Supplementary Materials.

### *In vivo* PK analysis

The mean POA exposure in plasma, ELF, and lung tissue over time following administration via the IV, PO, IT, or pulmonary routes are shown in **Supplementary Figure 4 A-C**. Plasma PK profiles of IV-POA (n=8) and IT-PM (n=4) are shown in **Supplementary Figure 4 A1**, while **Supplementary Figure 4 A2** depicts the plasma PK profiles following direct pulmonary administration of POA-formulations (PM (n=11), PML (n=10), and PLS (n=8)). PO POA achieved higher exposure, as the oral POA dose was the human equivalent dose, 300 mg/kg, which was much higher than that administered via the intravenous (120-fold), IT (150-fold IT-PM), or pulmonary (83-fold PM, 171-fold PML, and 162-fold PLS) routes (**Supplementary Figure 4** **A3**). The samples for characterizing POA PK in the ELF and lung tissue were sparse given the destructive nature of sampling. The PK profiles in the ELF and lung tissue are shown in **Supplementary Figures 4 B and 4 C**, respectively.

The observed NCA PK parameters (C_max_, T_max_, AUC_0-last_, AUC_total_, t_1/2_, and Mean residence time (MRT)) for POA in plasma for the different routes of administration and different formulations are reported in **Supplemental Table 1**. The half-lives for the different POA formulations following pulmonary administration (PM: 20.6 ± 6.95, PML: 20.3 ± 6.14, and PLS: 26.6 ± 17.3 min) and intratracheal PM (19.5 ± 14.8 min) were longer than the POA half-life in plasma following IV administration (12.5 ± 3.54 min) (**Supplemental Table 1 and Supplementary Figures 4 A1, A2**). The *T_max_* values following administration of the last pulmonary dose of PM, PML, and PLS were 31.1 ± 2.88, 37.5 ± 6.89, and 36.1 ± 10.9 min, respectively, delayed relative to IT administration of PM (17.7 ± 11.5 min) and attributable to slow release of POA from the inhalation site and the transit time to the absorption site in the lungs. Dose normalized POA exposure in the plasma following PO administration was significantly higher (**Supplementary Figure 4-A3**) (*AUC_total_* 1369 ± 481 µM·h (dose-normalized plasma *AUC_total_*: 11.4 µM·h)) than the pulmonary route (dose-normalized plasma *AUC_total_* for PM (1.15 µM·h), PML (2.15 µM·h), and PLS (1.89 µM·h)). The MRTs for PM, PML, and PLS in plasma were 47.9 ± 7.06, 52.9 ± 7.75, and 67.6 ± 184 min, while the IT-PM had an MRT of 33.6 ± 16.3 min. PO POA administration had the longest MRT of 143 ± 59.8 min.

Guinea pigs are a USDA protected species and the study was designed with an emphasis on the 3Rs (replacement, reduction, and refinement), sparse data was available for characterizing PK using NCA for ELF and lung tissue, which requires destructive sampling. For this reason, we utilized a PK/PD modeling approach to characterize the PK data and gain mechanistic insights into POA exposure in different parts of the lung.

### Mechanism-based PK model

A total of 187 plasma PK samples (IV: 37, IT: 11, PO: 60, and pulmonary: 79 (PM:37, PML: 24, and PLS: 18)) were collected from 50 guinea pigs, with 14/187 observations (7.5%) below the limit of quantification (BLQ, 202 nM). **Figure 3** depicts the schematic for the MBM developed to describe POA exposure measured in plasma, ELF, and lung tissue following IV, PO, IT, and pulmonary administration. POA PK was described using a three-compartment model with a linear clearance from plasma (CL_POA_) and volume of distribution in plasma (V_POA_). Linear intercompartmental clearance between lung and plasma (CL_d_) and between ELF and lung tissue (CL_ELF_) described POA distribution to the lung and ELF compartments. The transfer of POA from the inhalational depot to ELF was described by a first-order transfer rate, denoted by (*k_Form_*). The systemic bioavailability after inhalation was denoted by respective bioavailability terms (*F_Form_*), where “*Form*” represents each of the three POA dry powder inhalation formulations. To account for tissue partitioning of POA, a lung-to-plasma partition coefficient (*KP_Lung_*) was included in the model and was estimated separately for each POA inhalation formulation.

**Figure 3.**
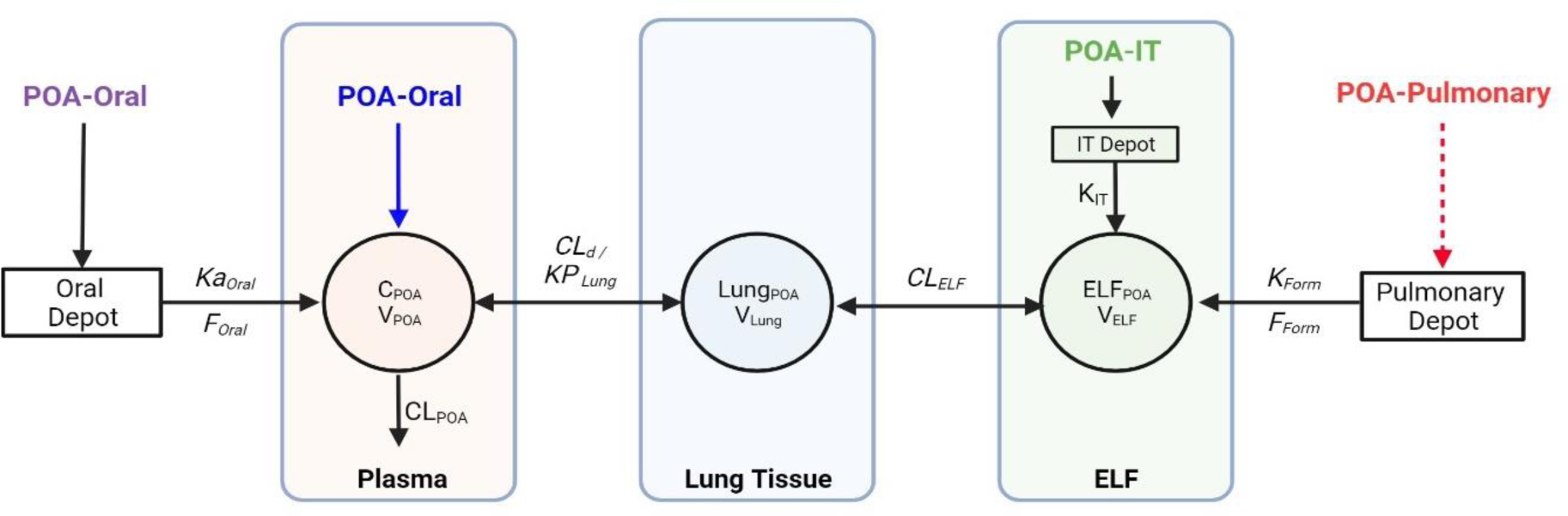
Mechanism-based pharmacokinetic (PK) model to describe the disposition of pyrazinoic acid (POA) in plasma, lung tissue, and ELF following per oral (PO), intravenous (IV), intratracheal (IT), and pulmonary administration of POA and POA formulations. For IV and PO PK studies, POA was dosed as a solution. For the IT PK study, PM powder was dosed. For pulmonary administration, spray-dried dry powder POA formulations were used. CLPOA, POA clearance; VPOA, POA central volume of distribution; CLd, POA distributional clearance between lung and plasma; V_lung_, volume of the lung tissue compartment; CLELF, POA distributional clearance between lung tissue and ELF compartment; VELF, volume of ELF compartment; *k_Form_*, first order transfer rate from pulmonary depot to ELF following pulmonary administration of POA formulations; *F_Form_*, bioavailability of POA formulations following pulmonary administration; *k_IT_*, first-order transfer rate to ELF from IT depot following IT administration of PM; *F_IT_*, bioavailability of intratracheally-administered PM; *Ka_Oral_*, first-order absorption rate following oral administration of POA; *F_Oral_*, bioavailability of orally administered POA; *KP_Lung_*, lung to plasma partition coefficient.^51^

A semi-physiological approach was adopted to describe the POA PK in ELF and lung tissue. The lung tissue volume (*V_Lung_*) was fixed to a literature-reported value of 2.0 x 10^-3^ L, which was consistent with the lung tissue volume calculated using the average body weight of the guinea pigs used in the study (2.1 x 10^-3^ ± 3.1 x 10^-4^ L).^49^ Similarly, guinea pig *V_ELF_* was fixed to the literature-reported value of 1 x10^-5^ L,^50^ consistent with the calculated *V_ELF_* of 3.86 x 10^-4^ ± 1.62 x 10^-4^ L (range: 4.71 x 10^-5^ − 8.56 x 10^-4^ L) using Equation 1. Clearance of POA from ELF, *CL_ELF_*, was estimated for each POA formulation, and depended on the dissolution of dry powders in ELF and release kinetics of POA from the different formulations. PK parameters *CL_d_*, *CL_ELF_*, *k_Form_*, *F_Form_*, and *KP_Lung_* for the systemic, intratracheal, and oral routes were fixed to the values estimated during the sequential development process. The *KP_Lung_* determines the partitioning of POA between the lung tissue and plasma, depending on the formulation.

The differential equations for simultaneously modeling the disposition of POA following administration of POA via the different routes of administration are described below.

#### IV POA and central compartment

The disposition of POA following IV administration is described by Equation (4):

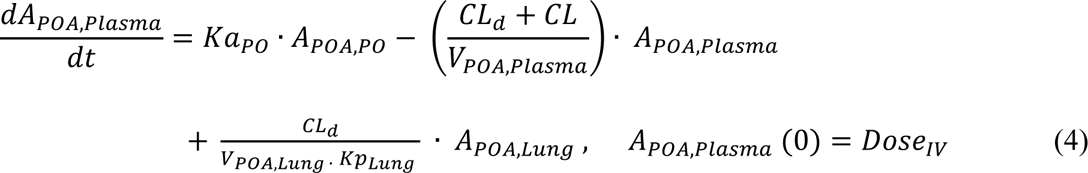

#### Oral POA

Disposition of oral POA is described by Equation (5):

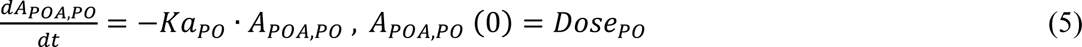

#### Pulmonary and IT dose of POA formulations

The pulmonary administration of POA formulations and intratracheal administration of PM via the depot compartment are described by Equations 6 and 7:

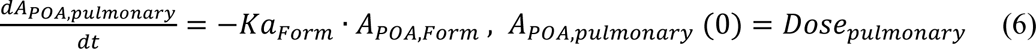

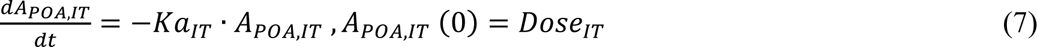

From the depot compartments, the POA distributes to the ELF compartment (Equation 8) followed by lung tissue (Equation 9) and plasma (Equation 4).

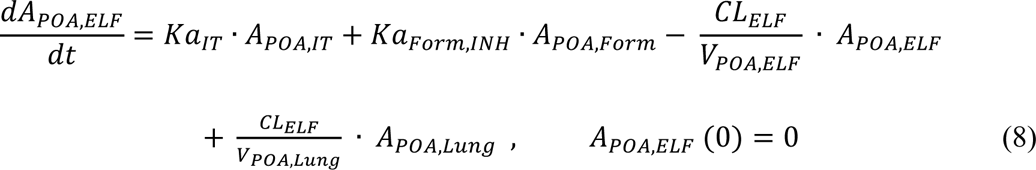

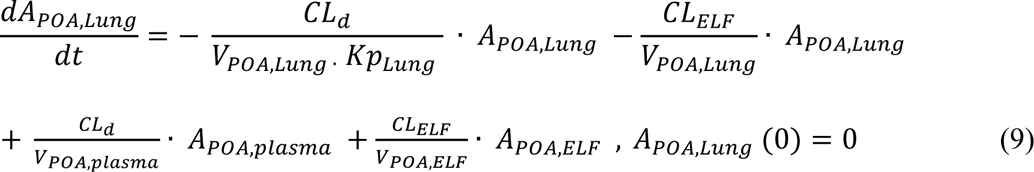

The residual unexplained variability is modeled using a combined additive plus proportional error model:

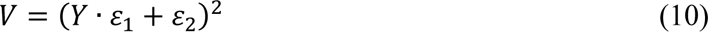

where V is the variability in the observed POA concentration, Y; ε_1_ (%), the proportional error, is based on the performance of the LC-MS/MS assay for quantifying POA (25%) in the different matrices, and ε_2_ (nM), an additive error, is fixed to half the lower limit of quantification (101 nM). The Beal3 method was used to handle the data BLQ.^52^

The model describes the observed PK data well for the different POA formulations and POA administered via different routes (**Figure 4 and Supplementary Figure 5**). Initially, a one-compartment model was developed to describe the observed IV PK with good precision. During subsequent model development, the systemic disposition parameters (CL_POA_ and V_POA_) were fixed to these estimated values. Similarly, oral POA PK was described by a one-compartment model with good precision, and the estimated absorption rate constant *Ka_Oral_* was estimated to be 0.958 h^-1^ and was fixed in the final model. Model performance was evaluated based on the goodness of fit plots (**Supplementary Figure 6-8**). The observed versus predicted POA concentration plots (**Supplementary Figure 6**) indicated that the model better captured POA PK for the IV (**Supplementary Figure 6A**) route and following pulmonary administration of PM (**Supplementary Figure 6D**) and PML (**Supplementary Figure 6E**) than POA administration via the oral (**Supplementary Figure 6B**) and IT (**Supplementary Figure 6C**) routes and for PLS (**Supplementary Figure 6F**) following pulmonary administration. The standardized residuals versus PRED and Time were evenly distributed around zero, indicating no major bias in the model. The final PK model parameters were estimated with good precision, with a CV% <32% (**Table 2**).

**Figure 4.**
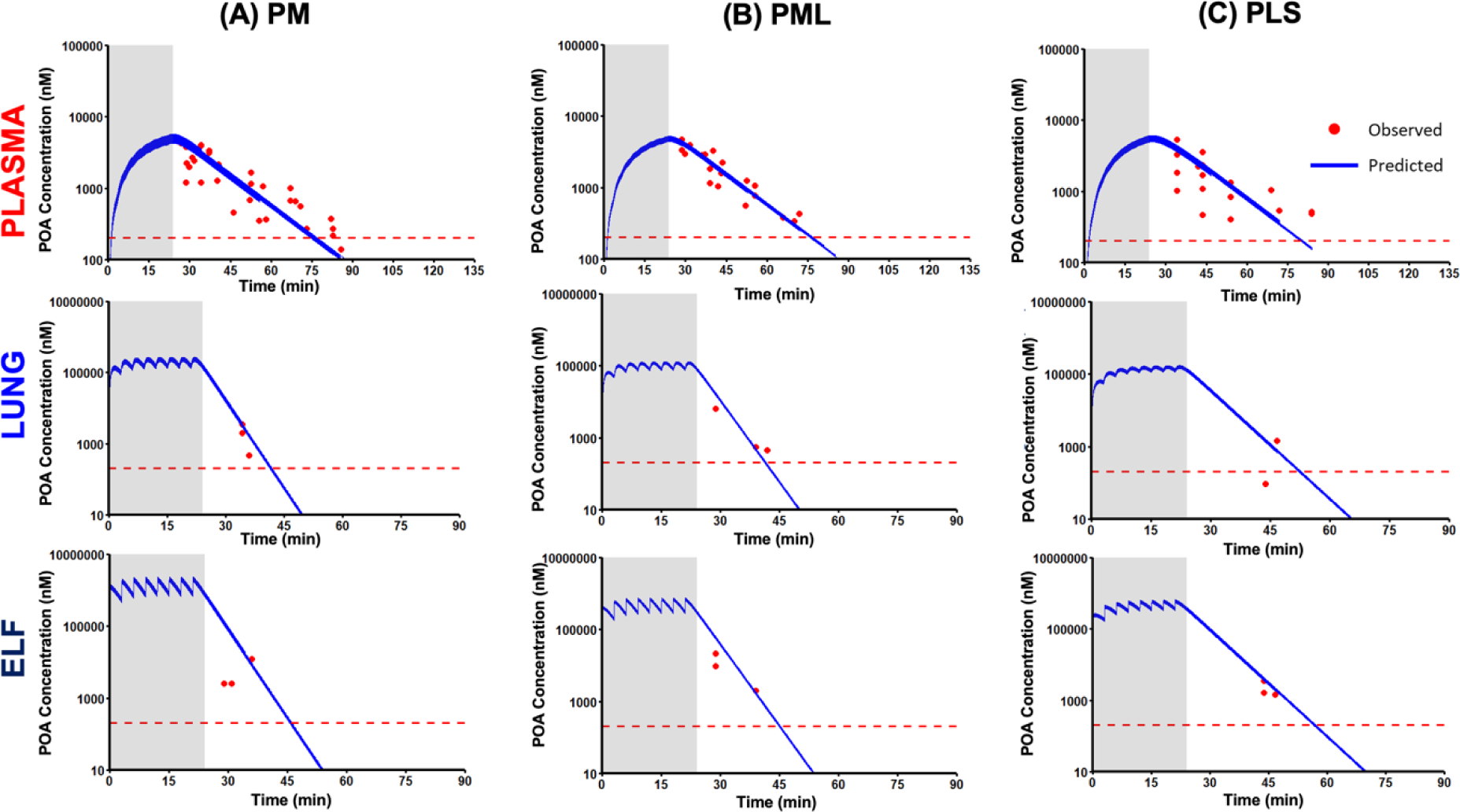
Mechanism-based model-predicted POA concentration versus time profiles following pulmonary administration of POA dry powder formulations **(A)** PM, **(B)** PML, and **(C)** PLS. Solid blue lines are model-predicted concentrations, while the red dots are the observed POA concentrations. The figure shows the predicted concentration versus time profiles in plasma (**top row**), lung tissue (**middle row**), and ELF (**bottom row**). Dashed red line indicates the lower limit of quantification (LLOQ 202 nM). Shaded region represents the total duration of 24 minutes over which the POA formulation was dosed.

**Table 2.**
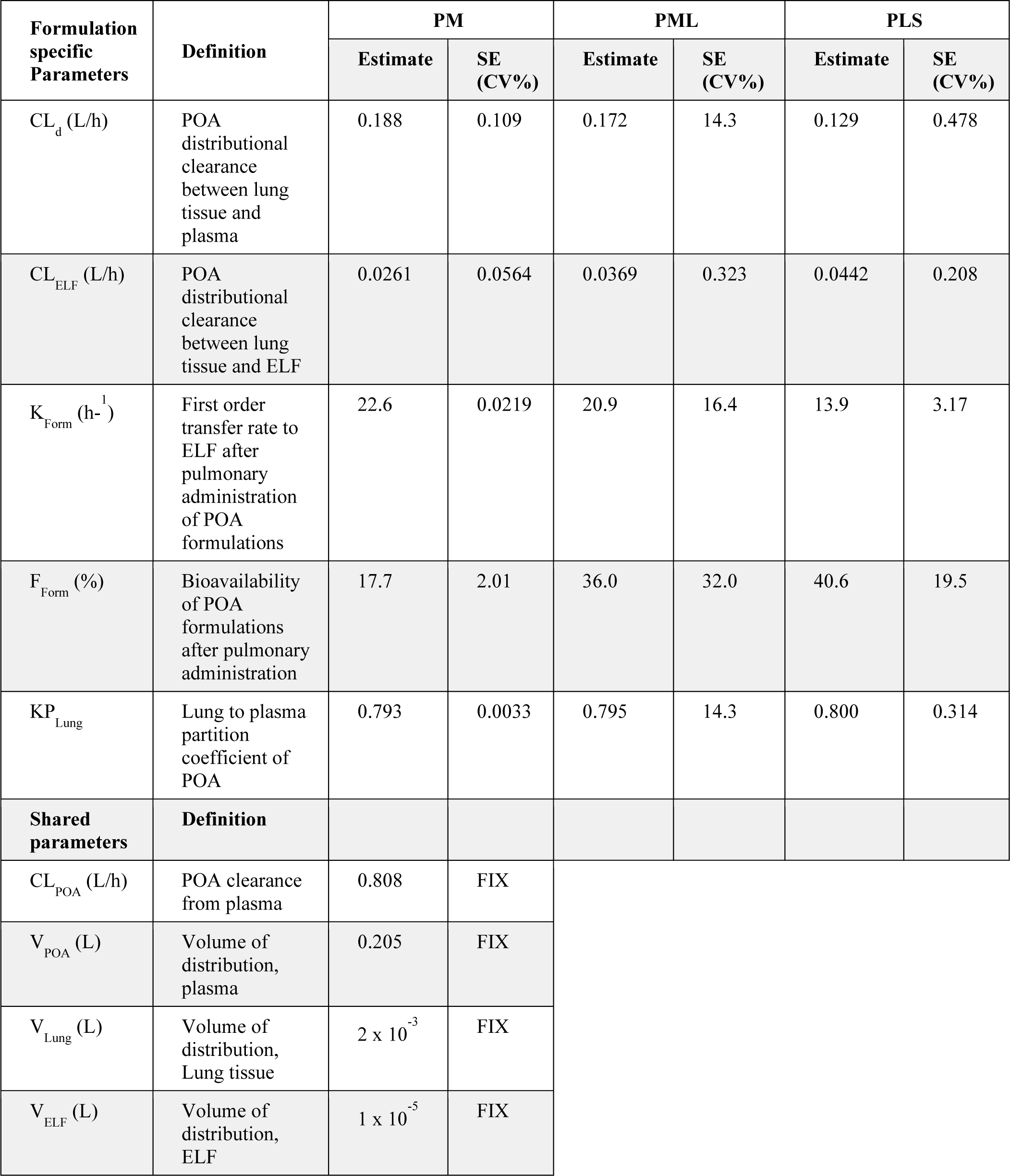
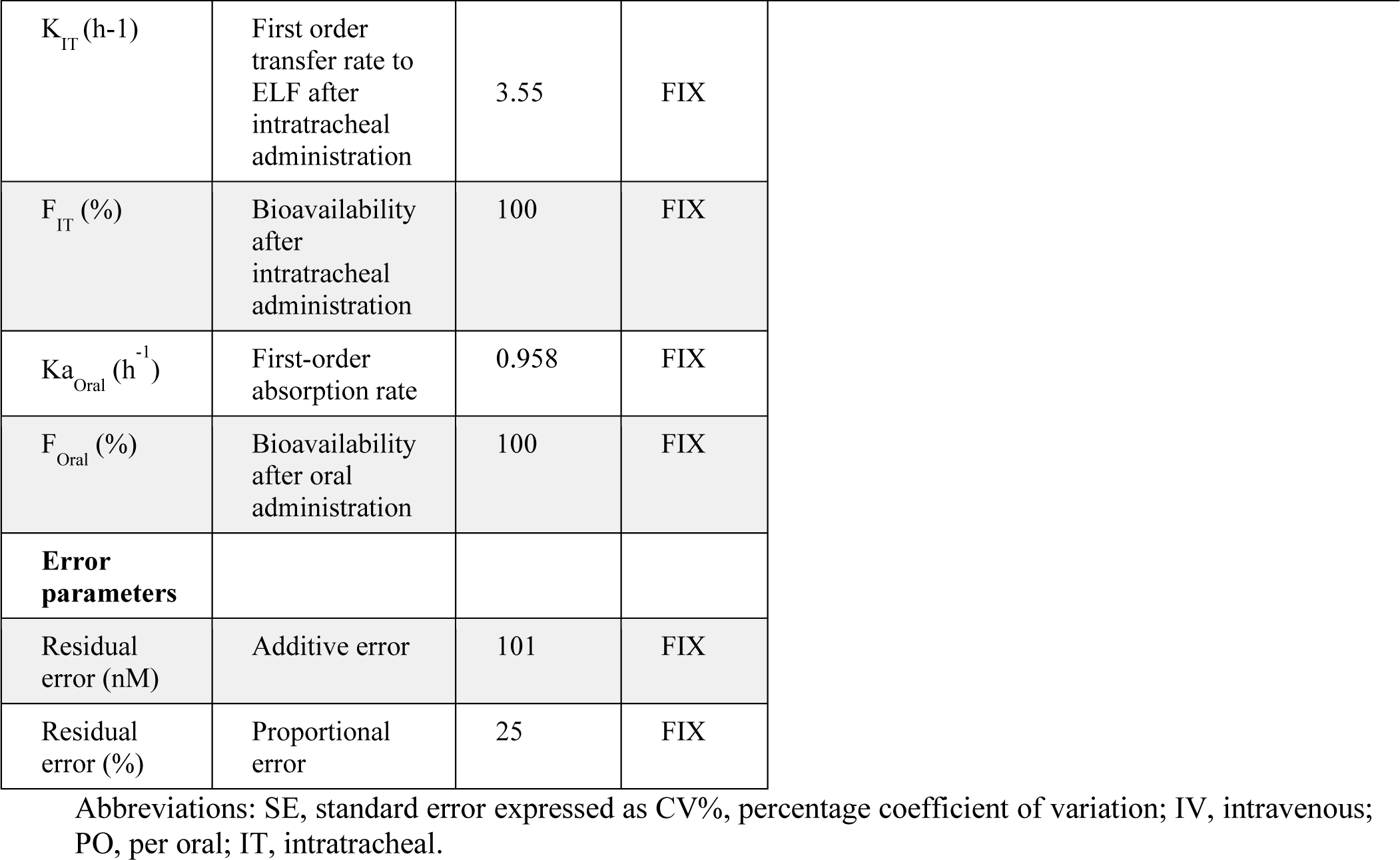
Mechanism-based PK model estimates for model parameters characterizing PK of POA in plasma, lung tissue and ELF compartments following PO, IV, IT and pulmonary administration of POA and POA dry powder formulations (PM, PML and PLS).

| Formulation specific Parameters | Definition | PM |  | PML |  | PLS |  |
| --- | --- | --- | --- | --- | --- | --- | --- |
|  |  | Estimate | SE (CV%) | Estimate | SE (CV%) | Estimate | SE (CV%) |
| $CL_d$ (L/h) | POA distributional clearance between lung tissue and plasma | 0.188 | 0.109 | 0.172 | 14.3 | 0.129 | 0.478 |
| $CL_{ELF}$ (L/h) | POA distributional clearance between lung tissue and ELF | 0.0261 | 0.0564 | 0.0369 | 0.323 | 0.0442 | 0.208 |
| $K_{Form}$ ($h^{-1}$) | First order transfer rate to ELF after pulmonary administration of POA formulations | 22.6 | 0.0219 | 20.9 | 16.4 | 13.9 | 3.17 |
| $F_{Form}$ (%) | Bioavailability of POA formulations after pulmonary administration | 17.7 | 2.01 | 36.0 | 32.0 | 40.6 | 19.5 |
| $KP_{Lung}$ | Lung to plasma partition coefficient of POA | 0.793 | 0.0033 | 0.795 | 14.3 | 0.800 | 0.314 |
| **Shared parameters** | **Definition** |  |  |  |  |  |  |
| $CL_{POA}$ (L/h) | POA clearance from plasma | 0.808 | FIX | | | | |
| $V_{POA}$ (L) | Volume of distribution, plasma | 0.205 | FIX | | | | |
| $V_{Lung}$ (L) | Volume of distribution, Lung tissue | $2 \times 10^{-3}$ | FIX | | | | |
| $V_{ELF}$ (L) | Volume of distribution, ELF | $1 \times 10^{-5}$ | FIX | | | | |

|  |  |  |  |
| --- | --- | --- | --- |
| $K_{IT}$ (h<sup>-1</sup>) | First order transfer rate to ELF after intratracheal administration | 3.55 | FIX |
| $F_{IT}$ (%) | Bioavailability after intratracheal administration | 100 | FIX |
| $Ka_{Oral}$ (h<sup>-1</sup>) | First-order absorption rate | 0.958 | FIX |
| $F_{Oral}$ (%) | Bioavailability after oral administration | 100 | FIX |
| **Error parameters** |  |  |  |
| Residual error (nM) | Additive error | 101 | FIX |
| Residual error (%) | Proportional error | 25 | FIX |

Abbreviations: SE, standard error expressed as CV%, percentage coefficient of variation; IV, intravenous; PO, per oral; IT, intratracheal.

The first order transfer rate to ELF from the pulmonary depot was higher for PM (*k_PM_*: 22.6 h^-1^) than PML (*K_PML_*: 20.9 h^-1^), while PLS had the slowest transfer rate (*K_PLS_*: 13.9 h^-1^). The model-predicted systemic bioavailability for PM (*F_PM_*:17.7%) was half that predicted for PML (*F_PML_*: 36.0%) and PLS (*F_PLS_*: 40.6%), indicating a lower systemic exposure for PM. The *KP_Lung_* was comparable for all three POA formulations (79%), indicating ∼79% of POA is available to partition between lung tissue and plasma.

The *AUC*_0−1.5*h*_, *C_max_*, and *T_max_* were calculated using the model-predicted plasma, ELF, and lung tissue concentrations over time profiles for the three POA formulations and for POA administered via the different routes (**Table 3**). The parameters reported in **Table 3** were dose normalized. Systemic exposure based on the dose-normalized *AUC*_0−1.5*h,Plasma*_ was higher for the IV (1.23 nM·h) and IT (1.25 nM·h) routes than the oral (0.840 nM·h) route. Of the POA formulations, PM had lowest systemic exposure (*AUC*_0−1.5*h,Plasma*_: 0.226 nM·h) compared with PML (0.455 nM·h) and PLS (0.503 nM·h).

**Table 3.** Comparison of model-predicted AUC, C_max_, and T_max_ of POA in plasma, lung tissue, and ELF compartments following PO, IV, IT, and pulmonary administration of POA and POA dry powder formulations (PM, PML and PLS).

| PK parameter | Route of administration | Formulation form | Plasma | Lung | ELF | Lung to plasma exposure ratio * | ELF to plasma exposure ratio * |
| --- | --- | --- | --- | --- | --- | --- | --- |
|  | **IV-POA** | Solution | 1.23 | 1.56 | 1.56 | 1.27 | 1.27 |
| **Dose normalized**<br>**AUC<sub>0-1.5h,pred</sub>**<br>**(nM·h)** | **PO-POA** | Solution | 0.840 | 1.05 | 1.05 | 1.25 | 1.25 |
|  | **IT-PM** | Solution | 1.25 | 6.83 | 45.1 | 5.48 | 36.2 |
|  | **Pulmonary-PM** | Dry powder | 0.226 | 6.87 | 46.0 | **30.4** | **203** |
|  | **Pulmonary-PML** | Dry powder | 0.455 | 7.35 | 34.4 | **16.2** | **75.7** |
|  | **Pulmonary-PLS** | Dry powder | 0.503 | 9.69 | 32.3 | **19.3** | **64.2** |
| **Dose normalized**<br>**C<sub>max,pred</sub>**<br>**(nM)** | **IV-POA** | Solution | 4.85 | 5.18 | 5.18 |  |  |
|  | **PO-POA** | Solution | 0.742 | 0.935 | 0.935 |  |  |
|  | **IT-PM** | Solution | 1.49 | 20.4 | 144 |  |  |
|  | **Pulmonary-PM** | Dry powder | 0.425 | 20.0 | 171 |  |  |
|  | **Pulmonary-PML** | Dry powder | 0.844 | 20.9 | 121 |  |  |
|  | **Pulmonary-PLS** | Dry powder | 0.908 | 25.8 | 98.2 |  |  |
| **T<sub>max</sub> (min)** | **IV-POA** | Solution | 0.091 | 2.48 | 2.52 |  |  |
| **T<sub>max</sub> (min)** | **PO-POA** | Solution | 28.8 | 29.4 | 29.4 |  |  |
|  | **IT-PM** | Solution | 13.4 | 2.58 | 1.06 |  |  |
|  | **Pulmonary-PM** | Dry powder | 23.9 | 21.6 | 15.5 |  |  |
|  | **Pulmonary-PML** | Dry powder | 24.1 | 22.0 | 18.5 |  |  |
|  | **Pulmonary-PLS** | Dry powder | 25.0 | 22.1 | 21.1 |  |  |

Abbreviations: IV, intravenous; PO, per oral; IT, intratracheal; C<sub>max,pred</sub> (nM), maximum predicted POA concentration; AUC<sub>0-1.5h,pred</sub> (nM·h), model-predicted area under the curve from 0 to 1.5 h for POA.
\* Exposure ratio = AUC<sub>Lung OR ELF</sub> / AUC<sub>Plasma</sub>

PM achieved a higher *C_max,ELF_* of 171 nM within 15.5 minutes of administration than PML (*C_max,ELF_* 121 nM; *T_max,ELF_*: 18.5 minutes) and PLS (*C_max,ELF_* 98.2 nM; *T_max,ELF_*: 21.1 minutes). This rapid *T_max,ELF_* for PM was correlated with the faster transfer rate (*k_PM_*: 22.6 h^-1^) from depot to the ELF compartment. PM had the highest ELF-to-plasma and lung tissue-to-plasma exposure ratios (ELF: 203; lung tissue: 30.4) compared with PML (ELF: 75.7; lung tissue: 16.2) and PLS (ELF: 64.2; lung tissue: 19.3).

Since PM is transferred at a higher rate into the ELF following administration, allowing it to achieve a higher peak concentration within a short timeframe and the ability to effectively penetrate both ELF and lung tissue, we regarded it as a superior formulation. PM had the lowest systemic bioavailability (i.e., lowest systemic exposure). Furthermore, PM achieved the highest exposure at likely sites of *Mtb* infection i.e., the AUCs in the ELF and lung tissue were high, which is necessary for efficacy of TB therapy and to limit off-target effects.

#### Scaling of human parameters and simulation of human dose

The developed mechanism-based PK model with published clinical PK data for PZA and POA were used to predict an efficacious human dose of inhaled PM dry powder. For scaling the human PK parameters, we considered the clearance and volume of POA from previously published clinical studies where PZA and POA PK were characterized after dosing of PZA.^39–41^ Other POA PK parameters estimated for PM in guinea pigs using our mechanism-based model were scaled using single species allometric scaling (**Supplemental Table 2**). We calculated the POA *C_max,ELF_* based on the ELF to plasma PZA concentration ratio and penetration of PZA to the ELF. Based on the literature reports the calculated POA *C_max,ELF_* ranges between 91.4 –134 mg/L.^20, 39, 44, 53, 54^ (**Supplemental Table 3**).

We performed our simulations targeting a *AUC_ELF_*/MIC similar to the above stated *AUC_Plasma_*/MIC target for PZA (9.68-14.5) considering a dose of 300 mg administered once a day (total dry powder 375 mg: 300 mg POA (80%) + 75 mg maltodextrin (20%)) or 150 mg administered twice a day (total dry power 187.5 mg: 150mg POA (80%) + 37.5 mg maltodextrin (20%)). The total daily dose achieved an *AUC_ELF_*/MIC of 14.1, and *C_max,ELF_* of 282 mg/L (range: 171 – 454 mg/L) for *Mtb* isolate with MIC of 8 mg/L (**Supplemental Table 4).** Given the limitation on the drug loading of the capsule in an inhalation device, the amount that can be delivered to the lung on a single breath (from a single capsule) and the number of capsules per dose, 187.5 mg administered twice a day is feasible. This calculated dose achieves the desired AUC and a *C_max_* > 35 µg/ml predictors of efficacy and would support human studies.

## DISCUSSION

Given the scarcity of new drugs effective against drug-resistant *Mtb*, new therapeutics would be a welcome addition to existing therapies. In addition, improved dosing strategies for existing TB drugs would help to ensure adequate exposure at the site of infection (i.e., ELF and lung tissue) resulting in improved bacterial clearance, shorten treatment duration, and will limit the emergence of resistance. PZA is an integral component of current TB therapy that can shorten treatment duration.^9, 10^ Among MDR-TB cases, defined as resistant to first-line drugs rifampicin and isoniazid, 60.5% are also resistant to PZA.^17^ The majority (∼90%) of PZA-resistant *Mtb* isolates have *pncA* mutations.^16^ To circumvent this most common category of PZA resistance, direct delivery of POA, the active moiety of PZA, is a promising approach. POA and POA esters have shown activity against PZA-resistant *pncA* mutant strains *in vitro*.^22, 37, 55^ However, when tested in mice or guinea pigs (unpublished data, Braunstein and Hickey), orally administered POA at the human-equivalent dose for PZA was not efficacious in either animal model, and higher oral POA doses are not feasible without significant tolerability problems (unpublished data, Braunstein and Hickey). ^21^

### The need for direct pulmonary administration of POA

Based on our analysis of oral and IV PK, the oral bioavailability was estimated to be 97-100%. Despite this high systemic exposure, oral POA had poor penetration into the ELF based on the NCA-calculated ELF-to-plasma and lung tissue-to-plasma exposure ratios of 0.15 and 0.65, respectively. These observations are consistent with lung and tuberculous lesion tissue-to-plasma ratios ranging between 0.2 and 0.6 for rabbits infected with *M. bovis* (a *pncA*-deficient strain) which were treated with PZA and POA was quantified.^20^ In a separate study, when POA was administered orally to *Mtb* infected C3HeB/FeJ mice at 450 mg/kg twice daily, the ELF-to-plasma exposure ratio was 0.45.^21^ This lower penetration into ELF after oral dosing may explain the lack of efficacy observed previously in mouse and guinea pig studies (unpublished data, Braunstein and Hickey).^21^

Monte Carlo simulations based on the relationship between drug exposure at the site of infection and the resulting killing activity obtained from *in vitro* hollow fiber infection^56^, pre-clinical, and clinical PZA PK data indicate that the drug exposure achieved in ELF is one predictor of long-term outcomes of PZA therapy. The sterilizing effect was associated with the PK/PD index, the ratio of exposure achieved in ELF (AUC_ELF_), and the minimum inhibitory concentration (MIC), a surrogate of PD response. Similarly, a C_max_ in plasma < 35 µg/ml was associated with PZA treatment failure and death.^47, 48^ The lack of efficacy of oral POA in mice may be explained by low exposure achieved in the ELF.^21^ To overcome these limitations, we previously evaluated nebulized POA esters, POA analogs that release POA independent of *Mtb pncA*. We also evaluated inhaled pyrazinoic acid/ester dry powder (PDP), a combination of POA and POA esters. Both inhaled POA esters and PDP were effective in combination with oral rifampicin for treating *Mtb-* infected guinea pigs compared with untreated animals. Moreover, the results indicated that inhaled PDP plus rifampicin was more effective than nebulized POA ester plus rifampicin, suggesting that the POA in the PDP dry powder formulation is more effective.^37, 55^

### POA dry powder formulations and impact of excipients on pulmonary PK

Here we developed three POA dry powder formulations for pulmonary delivery and evaluated their physicochemical characteristics and PK after direct pulmonary administration to guinea pigs. The aerodynamic diameter of dry powder particles and their flow and dispersion properties determine their ability to effectively penetrate deep into the lungs where *Mtb* reside.^34^ Maltodextrin and leucine were included as excipients to improve the flow characteristics of the dry powder formulations. Maltodextrin is a saccharide-based excipient with a lower dextrose equivalent (DE) rating and greater absorptive potential, as it exhibits a higher degree of branching and increased surface area. Its inclusion renders the formulation less hygroscopic while promoting its dissolution in mucus. Maltodextrin can provide a glassy matrix that helps improve stability and helps increase the encapsulation efficiency of the spray-dried powder.^57^

Leucine, when used with low solubility compounds such as azithromycin, showed accelerated dissolution attributed to increase in solubility of azithromycin.^58^ The solubility increase in azithromycin was dependent on the leucine concentration. Also, as leucine is a hydrophobic amino acid, it can potentiate the penetration of spray-dried powder across different cell membranes. This was evidenced by the high systemic exposure for PLS containing 50% leucine compared with PML containing 30% leucine.^58, 59^

Apart from improved dispersion, these excipients impacted the half-life of POA, helping to retain active POA and help redistribute it back to lung tissue and ELF to further contribute to efficacy. The plasma half-lives following pulmonary administration (IT: 19.5 mins; inhalation: PM: 20.6 mins, PML: 20.3 mins, and PLS: 26.6 mins) of POA dry powder formulations were longer than that after IV (12.5 mins) administration. Similar observations have been noted with capreomycin in humans, where its apparent half-life after pulmonary administration increased by ∼40% compared with its systemic half-life.^29, 31^ Similarly, in guinea pigs, the half-life of capreomycin increased by ∼36% with the addition of leucine (capreomycin: leucine 80:20).^29, 31^

### Pulmonary PK of POA and mechanism-based modeling

The inclusion of the innovative inhalational dosing strategy to ensure that the appropriate amount of active POA reached the site of infection limited our ability to obtain early PK samples (sampling was feasible after administration of the last dose, i.e., at 24 mins). This resulted in sparse PK samples for POA PK characterization in ELF and lung tissue, as a significant proportion of samples (64% in lung and 56% in ELF) were BLQ. Hence, our mechanism-based PK model was developed to effectively describe POA disposition following pulmonary administration, and this *in silico* model and its use to select a POA formulation achieving the highest exposure at the site of infection is the first of its kind.

Guinea pigs were chosen for this study as they are a well-established animal model for evaluating inhaled therapies, and they have specific advantages for evaluating anti-*Mtb* therapies because they develop hypoxic caseous granulomas with necrotic centers, the pathological hallmark of human TB.^60^ However, a challenge of working with guinea pigs, a USDA-protected species is the number of animals used to characterize PK in plasma, ELF, and lung tissue while adhering to 3Rs principles. This resulted in sparse PK samples for POA PK characterization in ELF and lung tissue, as a significant proportion of samples (64% in lung and 56% in ELF) were BLQ. Further, because the inhalational dosing strategy involved repeated doses administered every 3 minutes in the inhalation chamber this limited our ability to obtain early PK samples (sampling was feasible only after administration of the last dose, i.e., at 24 mins). Hence, our mechanism-based PK model was developed to effectively describe POA disposition following pulmonary administration to guinea pigs. The resulting *in silico* model and its use to select a POA formulation achieving the highest exposure at the site of infection is the first of its kind.

The approach used to develop this mechanism-based PK model was adopted from a model developed for nebulized colistin.^61^ Furthermore, using an innovative semi-physiological approach enabled us to optimize the model to estimate the POA exposure in ELF and lung tissue with improved precision. A similar approach has been used to describe the disposition of nebulized colistin in rats.^61^ Using this semi-physiological approach, we successfully estimated *k_Form_*, the model parameter describing the rate of transfer of POA to ELF after pulmonary administration of the POA formulation, to assist the selection of the optimal inhalation formulation. Of the three formulations evaluated here, the model predicted that PM had the fastest transfer rate to the ELF (22.6 h^-1^ vs. PML: 20.9 h^-1^ and PLS: 13.9 h^-1^). A lower MMAD is indicative of higher lung deposition,^62^ and PM had a comparatively lower MMAD than the other formulations. Based on transport by diffusion and mucociliary clearance in the lung, a fraction of POA will partition between the lung and plasma after pulmonary administration. Inclusion of partition coefficient *KP*_,*Lung*_ was necessary to account for this distribution in the lung. The estimated *KP*_,*Lung*_values were consistent across the three formulations, as POA can move from lung to plasma once it has reached the lung from the ELF, with POA transfer out of the lung independent of the excipients.

The model-based POA exposure predictions helped us to evaluate the penetration of these formulations into both ELF and lung tissue. The dose-normalized *AUC*_0−1.5*h,ELF*_ and *AUC*_0−1.5*h,Lung*_calculated based on model-predicted concentrations in the ELF and lung tissue compartments were higher for POA dry powder formulations than POA administered via the IV and oral routes. PM achieved the highest POA exposure in the ELF (*AUC*_0−1.5*h,ELF*_: 46.0 nM·h) and ELF-to-plasma exposure ratio (203) and the lowest systemic exposure (*AUC*_0−1.5*h,Plasma*_: 0.226 nM·h) of the inhaled POA formulations, indicating high pulmonary bioavailability. The low systemic exposure is attractive, as this will help to limit drug-related toxicity.,^21, 63^ as pulmonary POA will be used as part of multi-drug therapy with other antitubercular drugs.

### Feasibility of pulmonary POA and simulation of human dose

One of the important considerations for inhalation therapy with PM is the amount of powder that can be delivered safely. Capreomycin, at a dose of 300 mg, was administered to human participants in a phase I trial (360 mg total dry powder, 12 capsule of 30 mg)^31^. Commercially, a tobramycin dose of 112 mg is available for the treatment of cystic fibrosis and is administered via a Podhaler™ as 4 capsules, each containing 28 mg of tobramycin, twice daily. The inhaled dry powder dose achieved systemic exposure comparable to that achieved with 300 mg tobramycin inhalation solution (TIS, TOBI^®^, 300 mg tobramycin/5 mL).^64^ Colistin methanesulfonate (CMS) was delivered using a Twincer™ as a single dose^65, 66^ of 25-55 mg, while a CMS dose of 125 mg was administered twice daily using the EMA-approved Colobreathe_®_apparatus.^67^ In another phase I study, 32.5 mg of ciprofloxacin was delivered as a single capsule (50 mg total dry powder).^68^ For the different drugs administered via the inhalation route described above, the total dry powder delivered per day ranged from 250 – 360 mg, and no adverse events were reported except intermittent cough. Moreover, patient compliance and satisfaction were higher for dry powders than nebulization.

At present, no PK/PD index is established for POA, hence, based on the PK/PD index for PZA. Hence, the simulations to calculate the human dose for pulmonary administration targeted *AUC_ELF_*/MIC of 9.68-14.5 as the target PK/PD for pulmonary POA.^47, 48^ We also considered, plasma *C_max_* > 35 µg/ml as another predictor of efficacy.^48^ Using these predictors and anticipated limit on the amount of dry powder per dose, a dose of 150 mg POA (187.5 mg total PM dry powder (80% POA and 20% maltodextrin) given twice a day was calculated. At this dose the POA *C_max,ELF_* is above the calculated *C_max,ELF_* (91.4 –134 µg/ml) from clinical studies with *AUC_ELF_*/MIC of 14.1 for a *Mtb* strain with MIC of 8 µg/ml. Hence, the calculated PM dose (5.36 mg/kg per day or 2.68 mg/kg per dose) could achieve an efficacious *C_max_* and exposure at a much lower dose than clinical oral PZA regimen (15-30 mg/kg per day), leading to a reduction in systemic toxicities and preventing the development of bacterial resistance.

Our physicochemical characterization combined with *in vivo* and *in silico* PK analyses sets the stage for the future development of inhaled POA. However, the study design and approach have some limitations. Excipients counter strong interparticle cohesive forces and improve inspiratory flow rates and physiochemical characteristics. However, inclusion of excipients presents a challenge for balancing the percentage of excipient to total powder mass per dose to ensure that the total powder mass that can be inhaled in a single bolus or in rapid succession of actuations contains an adequate amount of drug, i.e., POA. This is compounded by the moderate lung delivery efficiencies that limit the maximum dose that can be delivered per dose via direct pulmonary administration without causing localized irritation or toxicity, which depends on the inspiration rate.^64^ PM consists of 80% POA and 20% maltodextrin, making it an ideal candidate with lower excipient and higher active drug content.

The results were obtained after inhalation of POA by guinea pigs in a custom-made chamber which do not represent the high efficiency of delivery that occurs with inhalers in humans. With the use of a dose calculation equation, we minimized non-specific losses of the administered dose when developing the model. Finally, there is no clinically established PK/PD index for POA, especially for its direct pulmonary administration. The currently available PK/PD index is based on the parent drug, PZA dosed orally, which might be misleading as the POA penetration into ELF is significantly lower following oral administration of PZA. Further PK/PD multiple daily-dose studies in guinea pigs infected with *Mtb* would help to further our knowledge about the inclusion of inhaled POA in the management and treatment of MDR-TB. Cytotoxicity studies to understand the relationship between POA exposure in the lung and peak concentration in the ELF would be beneficial in designing POA treatment regimens. In addition, further PK studies might be conducted in infected animals to evaluate the effect of disease on the disposition of drug.

## CONCLUSION

In conclusion, direct pulmonary administration of POA dry powder formulations achieved significantly higher exposure in ELF and lung tissue than POA administered via the oral and intravenous routes. Our mechanism-based model described the sparsely sampled data in lung and ELF. The data and model predictions suggest that PM is a suitable formulation for further characterization and evaluation in efficacy studies. PML has a PK profile comparable to PM and could serve as an alternate to PM. Future *in vivo* studies evaluating the POA exposure (PK) required for efficacy (PD) will further our knowledge about the inclusion of inhaled POA in the management and treatment of MDR-TB. The calculated human dose of 187.5 mg PM administered twice a day appears to be feasible via inhalation route (5.36 mg/kg for a 70 kg person) to achieve the necessary drug exposure associated with efficacy, which is encouraging for future clinical evaluation. This approach of using *in vivo* PK analyses combined with compartmental modeling will help future model-informed drug development (MIDD) of inhalational formulations for current and newer TB drugs.

## ASSOCIATED CONTENT

## AUTHOR CONTRIBUTIONS

MB, AH and GGR conceptualized and designed the research including the *in vivo* study and secured the funding. AH designed the POA formulations. IS formulated the spray-dried POA powders and performed physiochemical characterization. LR, CX, AA, AS, SY, VB conducted the *in vivo* PK analysis. RK, BJ, SY analyzed the *in vivo* data. SY, PH, and GGR modeled the data. SY, GGR, MB, and AH edited the manuscript.

## ACKNOWLEDGEMENTS

The authors acknowledge financial support from the National Institute of Allergy and Infectious Diseases through grant number R21AI131241 and the Potts Foundation.

## SUPPLEMENTARY INFORMATION

“ Supplementary information_Inhaled spray dried POA.docx”

**Supplementary Figure 1.** Experimental set-up for pulmonary dosing of guinea pigs. **(A)** Custom made plexiglass dosing chamber for dosing two guinea pigs simultaneously, **(B)** Custom made dosator with a female /male luer adapter to hold POA spray-dried powder formulations (PM, PML, and PLS) for administration of a single dose (8-10 mg).

**Supplementary Figure 2.** APSD from NGI for spray-dried POA (black), PM (blue), PML (red), and PLS (green) (bars appear in order from left to right). * Data is mean ± SD (n=3).

**Supplementary Figure 3.** X-ray powder diffraction: XRD pattern for, from bottom to top, POA (black squares), maltodextrin powder (red circles), L-leucine (blue triangles), and PML (pink stars).

**Supplementary Figure 4.** POA PK profiles in **(A)** plasma. **(A1)** Comparison of plasma POA PK following IV and IT administration of POA and PM, respectively, where IT administration results in slower rate of decline or longer half-life in plasma; **(A2)** Comparison of PK of POA formulations PM, PML, and PLS following pulmonary administration. There is no difference in POA half-life with the administration of the different excipients; **(A3)** POA PK in plasma following oral administration of POA compared with the pulmonary administration of POA formulations. POA PK following oral administration of a higher POA dose resulted in significantly higher exposure compared with pulmonary administration of a much lower POA dose, i.e., POA formulations. POA exposure for different POA formulations administered via the pulmonary route, intratracheal PM, and orally administered POA in the **(B)** ELF and **(C)** lung tissue. Dashed red line LLOQ (202 nM). Shaded regions in **(A2, A3)** represent the pulmonary administration period of 24 min.

**Supplementary Figure 5.** Predicted concentration versus time after **(A)** IV and **(B)** PO administration of POA and **(C)** IT-PM. Solid blue lines are model-predicted POA concentrations, and red dots are observed POA concentrations. The figure shows the predicted concentration versus time profiles in plasma (top row), lung tissue (middle row), and ELF (bottom row). Dashed red line indicates the lower limit of quantification (LLOQ 202 nM).

**Supplementary Figure 6.** Goodness-of-fit plots of the final mechanism-based PK model: observed versus population predictions for **(A)** IV-POA, **(B)** PO-POA, **(C)** IT-POA, **(D)** PM, **(E)** PML, and **(F)** PLS. The R^2^ values are as follows: IV: 0. 841; PO: 282; IT: 0.489; PM: 0.731; PML: 0.822; and PLS: 0.461.

**Supplementary Figure 7.** Goodness-of-fit plots for the final mechanism-based PK model: conditional weighted residuals versus population predictions (PRED) for POA administered as **(A)** IV, **(B)** PO, **(C)** IT-PM, and after pulmonary dosing of (**D**) PM, **(E)** PML, and (**F**) PLS. Dotted line indicates the line of unity. IV, intravenous; PO, per oral; IT, intratracheal.

**Supplementary Figure 8.** Goodness-of-fit plots for the final mechanism-based PK model: conditional weighted residuals versus time for POA administered as **(A)** IV, **(B)** PO, **(C)** IT-PM, and after pulmonary administration of (**D**) PM, **(E)** PML, and (**F**) PLS. Dotted line indicates the line of unity. IV, intravenous; PO, per oral; IT, intratracheal.

**Supplemental Table 1.** Plasma Non-compartmental analysis of POA and POA formulations following administration via the different routes

**Supplemental Table 2.** Scaling of human PK parameters for POA based on model predicted guinea pig parameters to estimate human dose of PM for pulmonary delivery.

**Supplemental Table 3.** Compilation of literature reported plasma and ELF Cmax values when PZA was dosed orally.

**Supplemental Table 4.** Simulated AUC and C_max_ after pulmonary delivery of PM at dose of 150 mg POA twice a day (187.5 mg PM twice a day) to 70 Kg human.

## ABBREVIATIONS

TB: tuberculosis
*Mtb*: *Mycobacterium tuberculosis*
PZA: pyrazinamide
POA: pyrazinoic acid
PK: pharmacokinetic
MBM: mechanism based pharmacokinetic model
ELF: epithelial lining fluid
MDR-TB: multi-drug resistant TB
DS-TB: drug-susceptible TB
PZAse: pyrazinamidase
PM: POA maltodextrin
PML: POA maltodextrin and leucine
PLS: POA leucine Salt
TGA: thermogravimetric analysis
NGI: Next Generation Impactor
XRPD: X-ray powder diffraction
APSD: aerodynamic particle size distribution
USP: United States Pharmacopeia
HPMC: hydroxypropyl methylcellulose
MMAD: mass median aerodynamic diameter
GSD: geometric standard deviation
FPD: fine particle dose
FPF: fine particle fraction
IACUC: Institutional Animal Care and Use Committee
JVC: Jugular vein catheter
BALF: bronchoalveolar lavage fluid
IV: intravenous
IT: intratracheal
PO: oral
NCA: Non-compartmental analysis
AUC: area under the concentration curve
t_1/2_: half-life
Tmax: time to peak concentration
Cmax: peak concentration
ML: maximum likelihood estimation
*Ka_Oral_*: first-order oral absorption rate
*F_Oral_*: oral bioavailability
*k_IT_*: first-order transfer rate between intratracheal depot to ELF
*F_IT_*: intratracheal bioavailability
*k_Form_*: first-order transfer rate of POA formulations between inhalation depot to ELF
*F_Form_*: bioavailability after pulmonary administration of POA formulations
BW_H_: body weight of humans
BW_GP_: body weight of guinea pig
P_H_: Scaled PK parameter for human
P_GP_: PK parameter for guinea pig
MRT: mean residence time
CL_POA_: POA clearance from plasma
V_POA_: volume of distribution in plasma
CL_d_: intercompartmental clearance between lung and plasma
CL_ELF_: intercompartmental clearance between ELF and lung
*KP_Lung_*: lung-to-plasma partition coefficient, *V_Lung_*, lung tissue volume
*V_ELF_*: volume of the ELF
MIC: minimum inhibitory concentration

