## SUPPLEMENTARY INFORMATION for "Pharmacokinetic considerations for optimizing inhaled spray-dried pyrazinoic acid formulations"

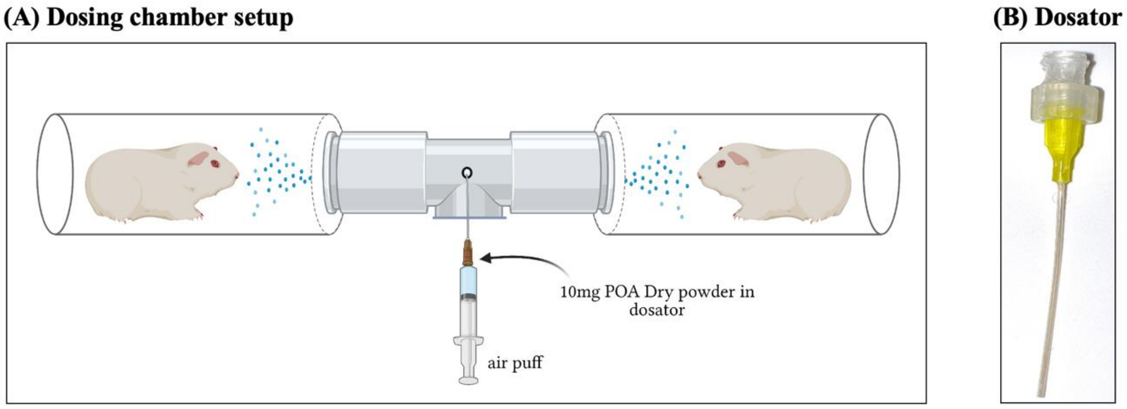


**Supplementary Figure 1.** Experimental set-up for pulmonary dosing of guinea pigs. **(A)** Custom made plexiglass dosing chamber for dosing two guinea pigs simultaneously, **(B)** Custom made dosator with a female /male luer adapter to hold POA spray-dried powder formulations (PM, PML, and PLS) for administration of a single dose (8-10 mg).

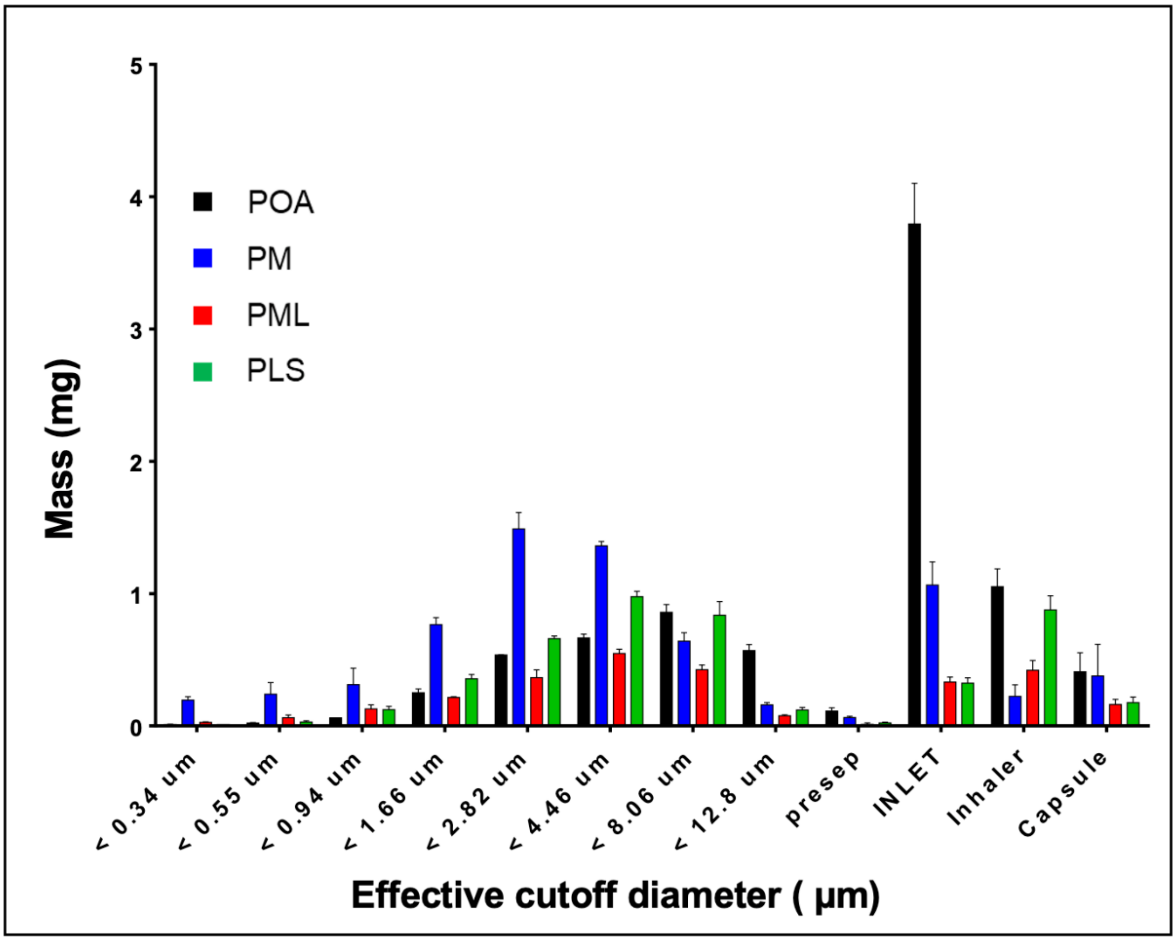


**Supplementary Figure 2.** APSD from NGI for spray-dried POA (black), PM (blue), PML (red), and PLS (green) (bars appear in order from left to right). * Data is mean ± SD (n= 3).

`

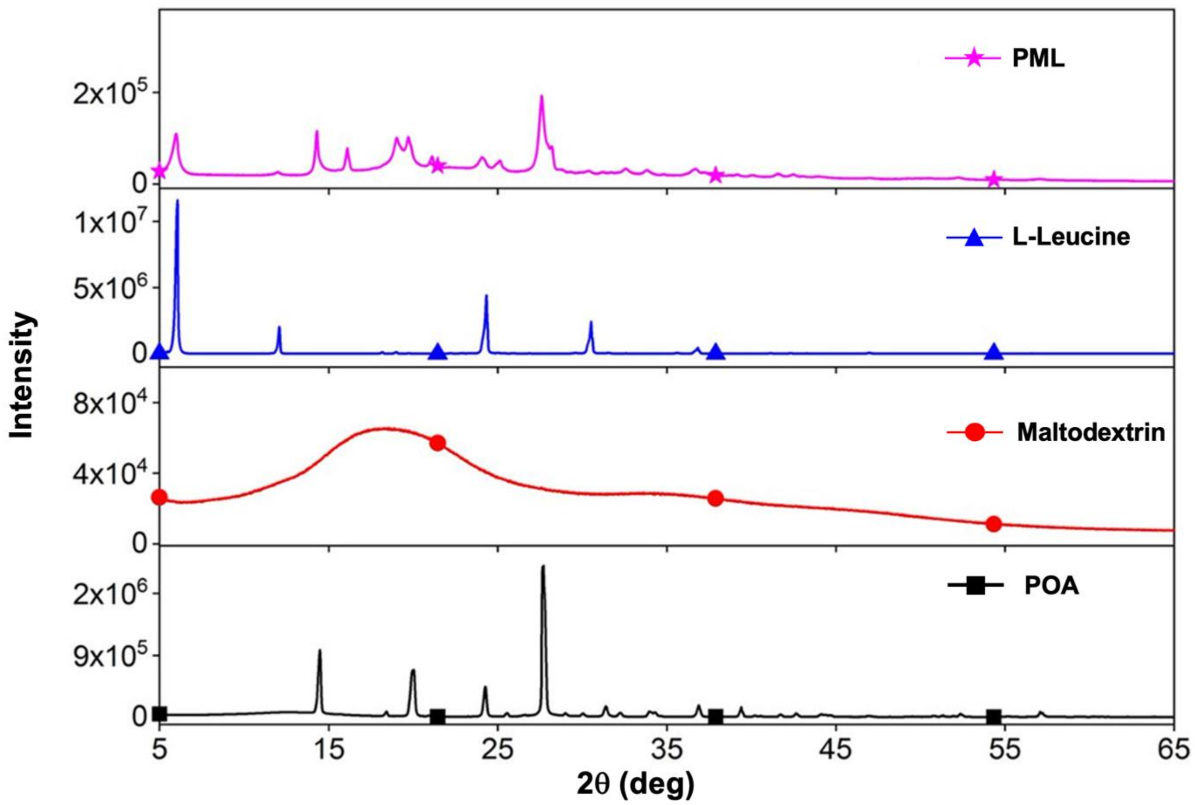


**Supplementary Figure 3.** X-ray powder diffraction: XRD pattern for, from bottom to top, POA (black squares), maltodextrin powder (red circles), L-leucine (blue triangles), and PML (pink stars).

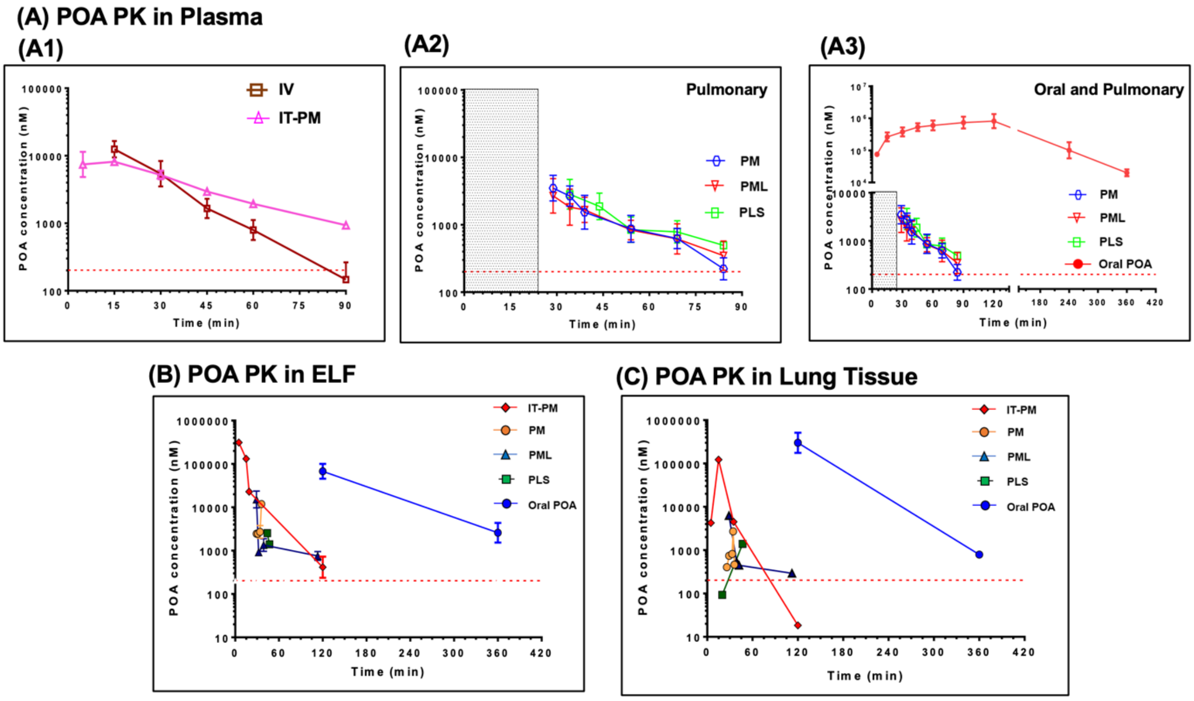


**Supplementary Figure 4.** POA PK profiles in **(A)** plasma. **(A1)** Comparison of plasma POA PK following IV and IT administration of POA and PM, respectively, where IT administration results in slower rate of decline or longer half-life in plasma; **(A2)** Comparison of PK of POA formulations PM, PML, and PLS following pulmonary administration. There is no difference in POA half-life with the administration of the different excipients; **(A3)** POA PK in plasma following oral administration of POA compared with the pulmonary administration of POA formulations. POA PK following oral administration of a higher POA dose resulted in significantly higher exposure compared with pulmonary administration of a much lower POA dose, i.e., POA formulations. POA exposure for different POA formulations administered via the pulmonary route, intratracheal PM, and orally administered POA in the **(B)** ELF and **(C)** lung tissue. Dashed red line LLOQ (202 nM). Shaded regions in **(A2, A3)** represent the pulmonary administration period of 24 min.

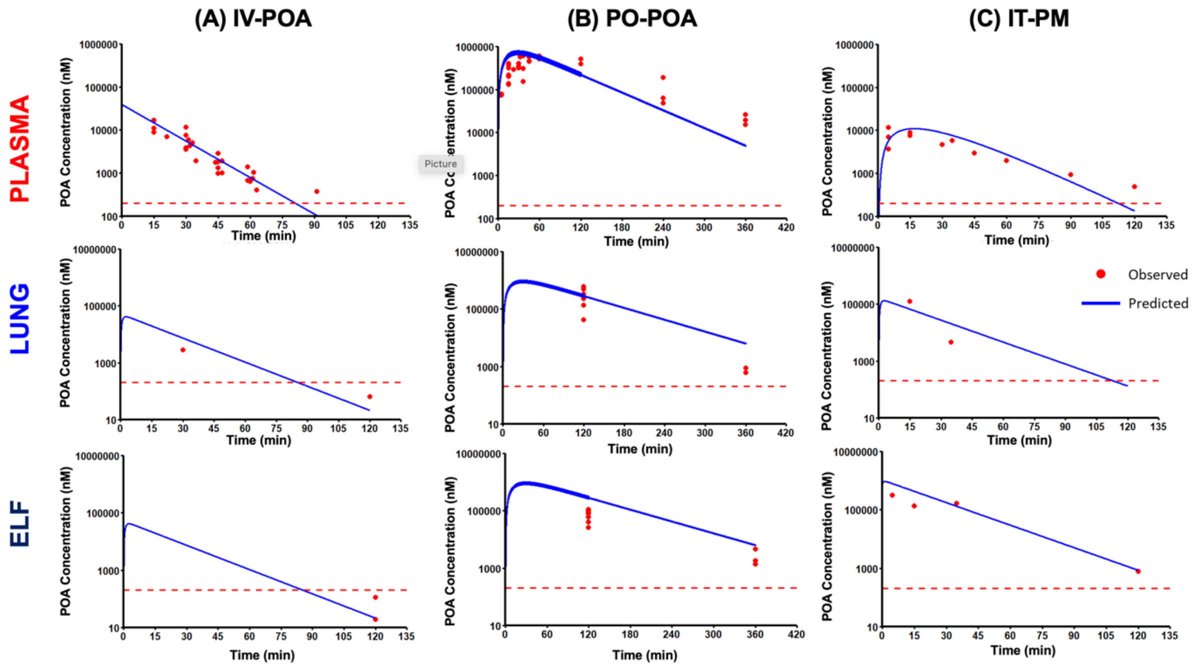


**Supplementary Figure 5.** Predicted concentration versus time after **(A)** IV and **(B)** PO administration of POA and **(C)** IT-PM. Solid blue lines are model-predicted POA concentrations, and red dots are observed POA concentrations. The figure shows the predicted concentration versus time profiles in plasma (top row), lung tissue (middle row), and ELF (bottom row). Dashed red line indicates the lower limit of quantification (LLOQ 202 nM).

`

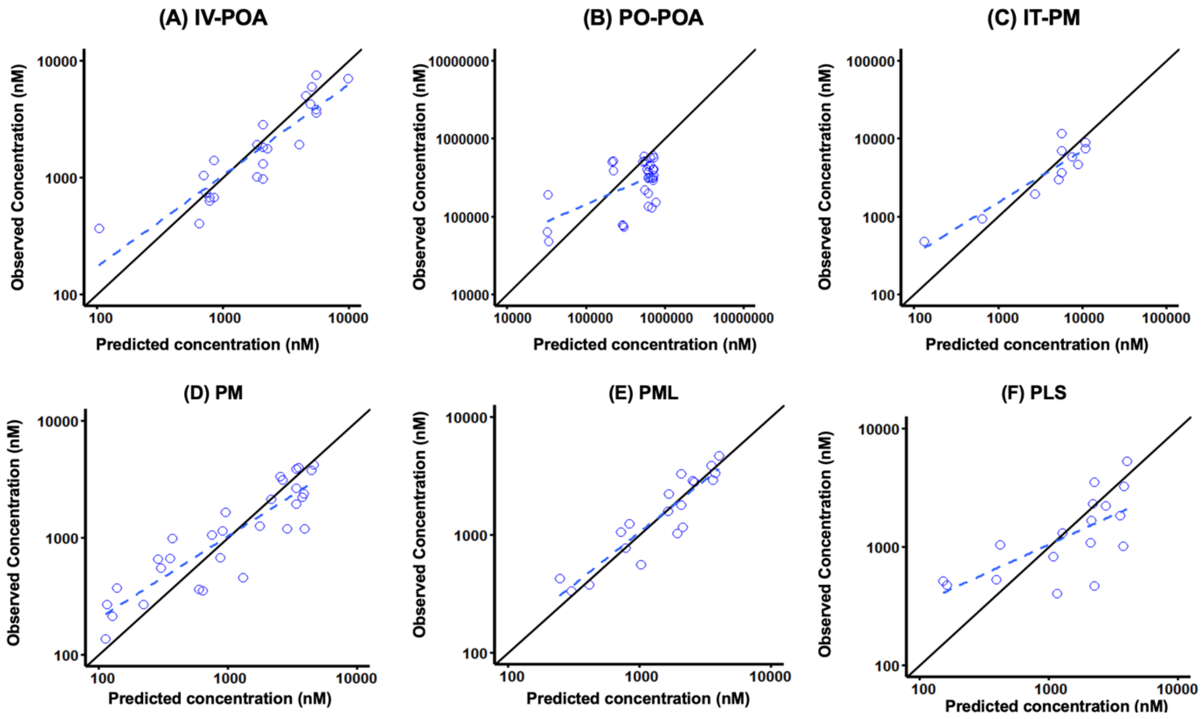


**Supplementary Figure 6.** Goodness-of-fit plots of the final mechanism-based PK model: observed versus population predictions for **(A)** IV-POA, **(B)** PO-POA, **(C)** IT-POA, **(D)** PM, **(E)** PML, and **(F)** PLS. The $R^2$ values are as follows: IV: 0. 841; PO: 282; IT: 0.489; PM: 0.731; PML: 0.822; and PLS: 0.461.

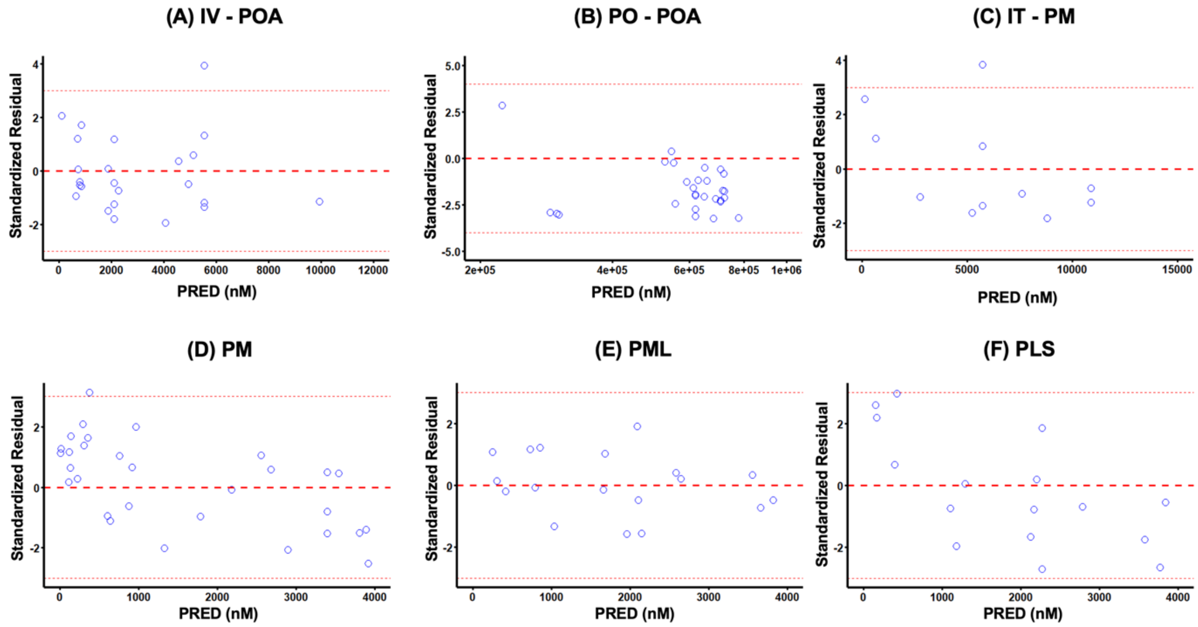


**Supplementary Figure 7.** Goodness-of-fit plots for the final mechanism-based PK model: conditional weighted residuals versus population predictions (PRED) for POA administered as **(A)** IV, **(B)** PO, **(C)** IT-PM, and after pulmonary dosing of (**D**) PM, **(E)** PML, and (**F**) PLS. Dotted line indicates the line of unity. IV, intravenous; PO, per oral; IT, intratracheal.

`

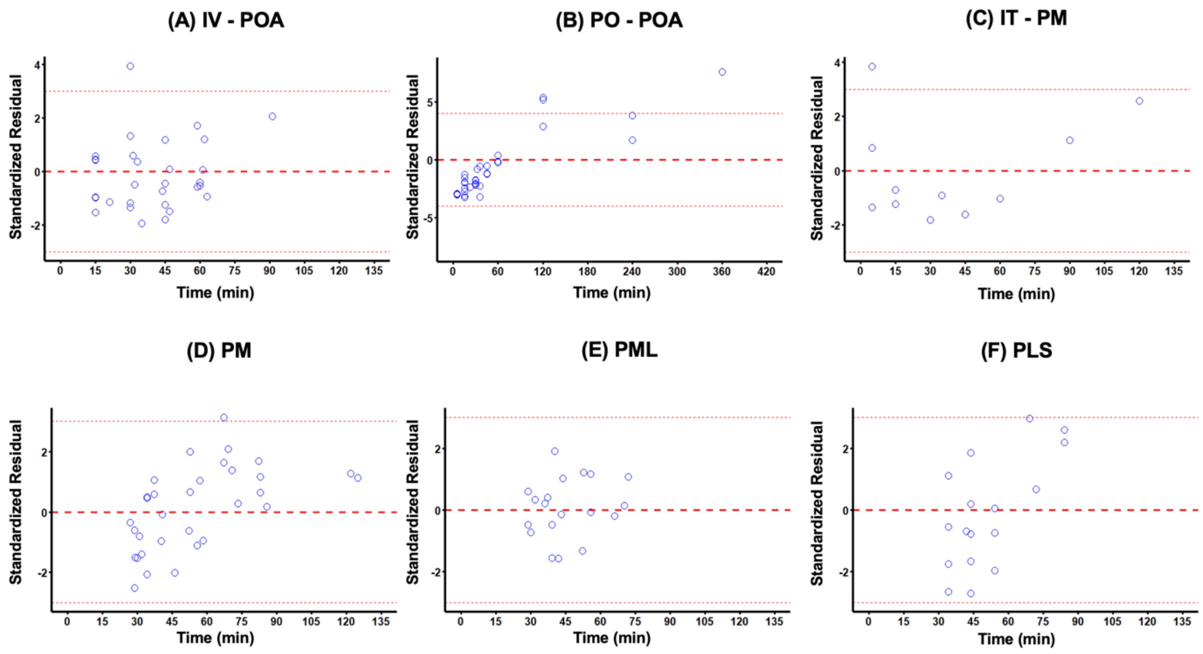


**Supplementary Figure 8.** Goodness-of-fit plots for the final mechanism-based PK model: conditional weighted residuals versus time for POA administered as **(A)** IV, **(B)** PO, **(C)** IT-PM, and after pulmonary administration of (**D**) PM, **(E)** PML, and (**F**) PLS. Dotted line indicates the line of unity. IV, intravenous; PO, per oral; IT, intratracheal.

**Supplemental Table 1.** Plasma Non-compartmental analysis of POA and POA formulations following administration via the different routes

| PK Parameters (Unit) | Definition | IV-POA | PO-POA | IT-PM | Pulmonary PM | Pulmonary PML | Pulmonary PLS |
|---|---|---|---|---|---|---|---|
| POA-Dose (mg) | Administered dose | 1.00 | 120 | 0.80 | 1.44 ± 0.134 | 0.702 ± 0.0365 | 0.737 ± 0.0522 |
| $C_{max}$ (μM) | Peak plasma concentration | 12.3 ± 3.72 | 857 ± 502 | 5.93 ± 3.94 | 2.82 ± 1.28 | 2.08 ± 1.56 | 3.21 ± 2.73 |
| $T_{max}$ (min) | Time to reach peak concentration | - | 88.7 ± 32.0 | 17.7 ± 11.5 | 31. 1 ± 2.88 | 37.5 ± 6.89 | 36.1 ± 10.9 |
| $T_{half}$ (min) | Half life of POA | 12.5 ± 3.54 | 52.8 ± 2.70 | 19.5 ±14.8 | 20.6 ± 6.95 | 20.3 ± 6.14 | 26.6 ± 17.3 |
| $AUC_{0\text{-}last}$ (μM·h) | AUC 0 – last | 8.87 ± 2.29 | 1032 ± 437 | 3.14 ± 2.10 | 1.46 ± 0.768 | 1.04 ± 0.779 | 1.54 ± 0.808 |
| $AUC_{total}$ (μM·h) | AUC total | 8.92 ± 2.32 | 1369 ± 481 | 5.88 | 1.65 ± 0.848 | 1.51 ± 0.848 | 1.76 ± 0.832 |
| Mean Residence Time (MRT) (min) | Mean residence time | 15.7 ± 1.56 | 143 ± 59.8 | 33.6 ± 13.6 | 47.9 ±7.06 | 52.9 ± 7.75 | 67.6 ± 18.4 |

The values are mean ± SD, Abbreviations : IV, intravenous; PO, per oral; IT, intratracheal

**Supplemental Table 2.** Scaling of human PK parameters for POA based on model predicted guinea pig parameters to estimate human dose of PM for pulmonary delivery.

| PK Parameter | Definition | Model Predicted Guinea Pig PK parameters | Human parameters | Comment |
|---|---|---|---|---|
| $CL_{POA}$ /F (L/h) | POA clearance from plasma | 0.808 | 13.4 | Published value [1,2,3] |
| $V_{POA}$ /F (L) | Volume of distribution, plasma | 0.205 | 51.4 | Published value [1,2,3] |
| $K_{Form}$ ($h^{-1}$) | First order transfer rate to ELF after pulmonary administration of PM | 22.6 | 6.29 | Allometric scaling [4] |
| $CL_d$ (L/h) | POA distributional clearance between lung tissue and plasma | 0.188 | 8.72 | Allometric scaling [4] |
| $CL_{ELF}$ (L/h) | POA distributional clearance between lung tissue and ELF | 0.0261 | 4.35 | Allometric scaling [4] |
| $KP_{Lung}$ | Lung to plasma partition coefficient of POA | 0.793 | 0.793 | Species independent |
| $V_{Lung}$ (L) | Volume of distribution, Lung tissue | 2 x 10-3 | 0.333 | Allometric scaling [4] |
| $V_{ELF}$ (L) | Volume of distribution, ELF | 1 x 10-5 | 0.002 | Allometric scaling [4] |

1. Mugabo et al, Eur J Drug Metab Pharmacokinet, 2019
2. Sundell J et al, AAC, 2021
3. Lacroix C et al, Eur. J. Clin. Pharmacol. 1989
4. Hanafin P et al, CPT-PSP, 2021

**Supplemental Table 3.** Compilation of literature reported plasma and ELF Cmax values when PZA was dosed orally.

| | Mugabo et al, 2019 [4] | Via et al, 2015 [5] | Naftalin et al, 2017 [2] |
|---|---|---|---|
| **Human Dose (mg/kg)** | 25 | 22-30 | 25 |
| **Patient population** | TB With and Without HIV | TB + HIV negative | TB + HIV negative |
| **Plasma Cmax (mg/L)** | | | |
| **PZA** | 25.8 (5.88–53.1) | 43.0 | 34.8 (31.9–41.9) |
| **POA** | 6.7 (1.50–20.0) | 7.0 | 4.57 (3.2–6.2) |
| **PZA/POA Cmax ratio in Plasma** | 3.85 | 6.14 | 7.62 |
| **PZA ELF penetration is 17.8- fold @ 2h and 22-fold @ 4h: Considering an average 20-fold penetration of PZA into ELF [1,3]** | | | |
| **Calculated PZA: Cmax in ELF** | 516 | 860 | 696 |
| **Observed ELF concentration of PZA 431 ± 220 mg/L[1]** | | | |
| **Calculated POA : Cmax in ELF based on PZA/POA Cmax ratio in plasma** | 134 | 140 | 91.4 |

* TB- Tuberculosis

**Supplemental Table 4.** Simulated AUC and $C_{max}$ after pulmonary delivery of PM at dose of 150 mg POA twice a day (187.5 mg PM twice a day) to 70 Kg human.

| PK Parameter | Definition | 150 mg BiD Median (Min –Max) |
|---|---|---|
| $AUC_{ELF}$ (mg·h/L) | AUC of the ELF after pulmonary delivery | 113 |
| $AUC_{ELF}$ /$MIC_{(8\ mg/L)}$ | AUC above MIC for *Mtb* isolate with MIC of 8 mg/L | 14.1 |
| $AUC_{Lung\ tissue}$ (mg·h/L) | AUC of the lung tissue after pulmonary delivery | 43.5 |
| $AUC_{Plasma}$ (mg·h/L) | AUC of the plasma after pulmonary delivery | 21.8 |
| $Cmax_{ELF}$ (mg/L) | Peak concentration in ELF after pulmonary delivery | 282 (171 – 454 ) |
| $Cmax_{Lung\ tissue}$ (mg/L) | Peak concentration in lung tissue after pulmonary delivery | 96.1 (56.5 – 173) |
| $Cmax_{Plasma}$ (mg/L) | Peak concentration in plasma after pulmonary delivery | 2.92 (1.37 – 4.85) |

### SUPPLEMENTARY METHODS

#### Manufacturing of PM, PML, and PLS dry powders

Briefly, POA and excipients (maltodextrin and leucine) were dissolved in 100% ultrapure water (5 mg/mL w/v solids for POA and PM) or 4:1 water: EtOH (v/v) (10 mg/mL w/v solids for PML). The resulting solutions were fed via a peristaltic pump though a heated nozzle and under atomizing nitrogen flow ($N_2$) to rapidly evaporate the solvent, forming dry powder particles. All powders were collected downstream via a cyclone (standard Buchi B290 cyclone) in amber vials and stored at room temperature with desiccant. Samples were aspirated with dehumidified air directed into the opening (22-27°C and 22-35% RH). For manufacturing POA and PM (4:1 mass ratio) dry powders, aspiration, inlet temperature, peristaltic pump speed, and $N_2$ were set to 100%, 120°C, 2 mL/min (5%), and 667 L/h (40 mm on rotameter), respectively. The resulting outlet temperatures were 55-60°C. For manufacturing PML (4:3:3 mass ratio) dry powder, aspiration, inlet temperature, peristaltic pump speed, and $N_2$ were set to 100%, 85°C, 3 mL/min (10%), and 1052 L/h (50 mm on rotameter), respectively. Resulting outlet temperatures were 40-45°C. PLS was synthesized as previously reported. The dry powder was generated with a Buchi B290 spray dryer using the standard cyclone. The PLS was dissolved in ultrapure water (> 17 MΩ) at a concentration of 10 mg/mL. The aspirator, inlet temperature, pump speed, and $N_2$ were set to 35 $m^3/h$, 140°C, 6.5 mL/min, and 1052 L/h (50 mm on rotameter), respectively. Resulting outlet temperatures were 60-70°C. All powders were collected and stored under desiccant.[1]

#### Physicochemical characterization

*Dry powder morphology*

Dry powder morphology was assessed by scanning electron microscopy (SEM; Quanta 200 SEM (FEI, Hillsborough, OR, US)). A small amount of powder was deposited on aluminum stubs topped with double-sided copper tape. Samples were sputter coated with Au/Pd (Hummer Sputtering System, Anatech Ltd., Union City, CA, US) for 2 min before SEM images were taken. Imaging was performed using an accelerating voltage of 5-10 kV and spot size of 3-3.5.

*Thermal analysis*

Thermal analysis and moisture content were determined by thermogravimetric analysis (TGA; Q50 TGA, TA Instruments, New Castle, DE, US). About 5-10 mg of powder were loaded in platinum TGA sample holders. Under an $N_2$ gas stream, data were collected over a temperature range of 30°C – 500°C using a 5°C/min ramp rate.

*Aerodynamic characterization*

A nominal powder mass of 10 mg (including excipients) was loaded into a #3 hydroxypropyl methylcellulose (HPMC) capsule and placed in a RS00 Mod.8 inhaler (Plastiape, Osnago, Italy). The inhaler was actuated and placed in a mouthpiece adapter of the NGI inlet under vacuum pressure. NGI stages and inlet, the RS00, and capsule were washed with a known volume of deionized water to collect the deposited mass. Samples were assayed at 269 nm via UV-Vis spectroscopy (SynergyMX, Biotech, Winooski, VT, US) for POA content to construct Aerodynamic Particle Size Distributions (APSDs). Masses collected at each stage were plotted against the cumulative percent undersize, which was then used to plot stage cutoff diameter vs the corresponding probability scale. The latter was used to calculate the mass median aerodynamic diameter (MMAD) by fitting a log-linear line to the two points on either side of the 50% cumulative mass to determine the median. The geometric standard deviation (GSD) was determined by calculating the square root of the ratio of particle size at the 84th and 16th percentiles of the particle size distribution. Fine particle dose (FPD) was determined by summing the masses of POA deposited at stage 3 and below based on NGI (particle sizes < 4.46 µm).

`

*Crystallinity*

The powders were characterized for crystallinity by X-ray powder diffraction (XRPD) using a D8 instrument from Bruker AXS Inc. (Karlsruhe, Germany). A solid mass was transferred to a flat, low diffraction silicon wafer and scans were performed from 5 – 65° 2θ at intervals of 0.02° 2θ with a 2 s dwell time at 40 kV and 40 mA using a copper anode beam source (0.154 nm wavelength). Jade version 9.6 software (Materials Dara Inc. Livermore, CA, US) was used to process and analyze resulting patterns.

**Inhalational and intratracheal pharmacokinetics**

*Dosing chamber and dosator for inhalational dosing*

A custom-built plexiglass, nose-only guinea pig aerosol dosing chamber (**Supplemental Figure 1A**) was used to perform inhalation studies. Two guinea pigs were dosed simultaneously, and the powder was dispensed using a custom-made single use dosator (**Supplemental Figure 1B**) developed previously for dry powder delivery.[2] A plastic (PTFE) luer needle (20 G; McMaster-Carr, Elmhurst, IL) was employed for ease of powder delivery rather instead of a metal needle. A female/male adapter luer adapter was used to hold the dry powder with a small stainless steel mesh (SST 304; 42 X 42 X 0.0055 from McMaster-Carr) punched using a die to achieve a final outer diameter of 5 mm. Once the powder was added, parafilm was used to cover the dosators while they were stored with desiccant prior to use. Three to five actuations of 2 mL of air were used to ensure that all the powder was delivered into the animal dosing chamber while only one actuation of 2 mL was used during direct insufflation.

*Calculation of actual inhaled dose*

The actual inhaled dose was calculated based on a combination of physiological parameters established for guinea pig and parameters established for the dosing chamber using the relationship described in the equation below:[3]

$$Actual\ inhaled\ dose(mg) = \frac{Drug\ in\ dosator\ [mg] \times duration\ of\ dosing\ [min] \times breathing\ rate\ \left[\frac{breaths}{min}\right] \times tidal\ volume\ \left[\frac{L}{breath}\right] \times Lung\ fraction[\%]}{Exposure\ chamber\ volume\ [L] \times TurnoverNumber}$$

1. Breaths/min are calculated based on relation[4]: $295/Body\ weight^{0.25}$
2. Tidal volume was computed[4] as: $0.0074\ \times Body\ weight\ Cm^{3}$
3. Lung fraction was set to 6% based on the MMADs for all aerosols being in the range 2.5-3.0 µm and GSDs of 1.6-1.7[4]
4. Exposure chamber volume: 0.83 L

### SUPPLEMENTARY RESULTS

#### Aerodynamic particle size distribution data interpretation

The quantities of POA in the samples collected at each stage were plotted against the cumulative percent undersize, which was then used to plot the effective cutoff diameter vs the corresponding probability scale. Each stage of the NGI is calibrated to collect particles of known aerodynamic particle size. The collection efficiency at each stage is defined by the cut-off aerodynamic particle diameter above which particles are collected and below which they pass to the next stage. The mass of drug collected at each stage is used to reconstruct the particle size distribution as the cumulative percentage under the cut-off size for each stage. The aerodynamic particle size distribution is plotted as the cumulative percentage undersize, by mass, on a probability scale against the logarithms of cut-off aerodynamic particle diameters. Data from this plot were used to calculate the mass median aerodynamic diameter (MMAD) by fitting a log-linear line to the two points on either side of the 50% cumulative mass to determine the median. Geometric standard deviation (GSD) was determined from the square root of the ratio of particle sizes at the 84th percentile and the 16th percentile of the particle size distribution.

**Supplementary Figure 2** shows the mass deposited at each stage of the NGI (y-axis) as a function of calibrated cut off aerodynamic particle diameter (x-axis). POA had a higher median diameter, as indicated by the peak in the distribution, compared with the other formulations. In addition, POA did not disperse as an aerosol as efficiently as indicated by deposition in the NGI inlet. The statistically-derived parameters of MMAD and GSD that specify the central tendency and breadth, respectively, of the aerodynamic particle size distribution are given in **Table 1**.

**X-Ray powder diffraction**

The particles were characterized for crystallinity by X-ray powder diffraction (XRPD, Model D8 instrument, Bruker AXS Inc., Karlsruhe, Germany).

**Supplementary Figure 3** shows XRP diffractograms for POA, PLS, and PLM. XRPD demonstrated that POA, LEU, and PLM exhibited crystallinity, the latter showing slight amorphous qualities, likely due to the maltodextrin content, which was amorphous as-received. Spray-dried PLS has previously been shown to be amorphous in structure.[1]

**Model evaluation**

The mechanism-based model-based fits for POA in plasma (top row), lung (middle row), and ELF (bottom row) for IV POA (A), PO-POA (B), and IT-PM (C) are shown in **Supplementary Figure 5**. Model evaluation was based on i) observed versus predicted concentrations via the IV, PO, and IT routes (**Supplementary Figure 6 A-C**) and for PM, PML, and PLS (**Supplementary Figure 6 D-F**); ii) conditional weighted residuals versus PRED for the IV, PO, and IT routes (**Supplementary Figure 7 A-C**) and for PM, PML, and PLS

`

(**Supplementary Figure 7 D-F**); and iii) standardized residuals over time for the IV, PO, and IT routes (Supplementary **Figure 8 A-C**) and for PM, PML, and PLS (Supplementary **Figure 8 D-F**).

**Simulation of human dose**

**Supplemental Table 2** shows the scaled human PK parameters for pulmonary POA. Simulations were performed using a human dose of 150 mg POA administered twice a day (total dry powder 187.5 mg of PM) using scaled PK parameters. **Supplemental Table 3** shows the literature review of the plasma and ELF $C_{max}$ data for PZA, which was used to estimate the $C_{max}$ of POA in the ELF. The calculated human parameters (AUC and $C_{max}$) obtained using the human dose of 150 mg POA are reported in **Supplemental Table 4.**
